# Leveraging Natural Language Processing models to decode the dark proteome across the Animal Tree of Life

**DOI:** 10.1101/2024.02.28.582465

**Authors:** Gemma I. Martínez-Redondo, Francisco M. Perez-Canales, José M. Fernández, Israel Barrios-Núñez, Marçal Vázquez-Valls, Ildefonso Cases, Ana M. Rojas, Rosa Fernández

## Abstract

Functional annotation is crucial in biology, but many protein-coding genes remain uncharacterized, especially in non-model organisms. FANTASIA (Functional ANnoTAtion based on embedding space SImilArity) integrates protein language models for large-scale functional annotation. Applied to ∼1,000 animal proteomes, it predicts functions to virtually all proteins, revealing previously uncharacterized functions that enhance our understanding of molecular evolution. FANTASIA is available on GitHub at https://github.com/CBBIO/FANTASIA.

## Main

Understanding protein function is essential for deciphering genome evolution and the complexity of life. Gene Ontology (GO) provides a standardized vocabulary to describe gene functions across species^1^, organizing terms into three categories: biological process, molecular function, and cellular component. This framework is crucial for interpreting genome-wide data, yet a substantial portion of protein-coding genes, particularly in non-model organisms, but also in model organisms to a lesser extent, remain unannotated—forming what we call the “dark proteome”.

Traditional functional annotation methods^2,3^ rely on sequence homology, limiting their ability to annotate highly divergent proteins. Many genes evade detection due to rapid evolution, orphan status, or intrinsic disorder, restricting their functional classification. For instance, ∼30% of proteins in the model *Caenorhabditis elegans* lack a function in Uniprot^4^, while 41% and 50% of genes in a tardigrade^5^ and sponge^6^ species, respectively, remained unannotated. As large-scale genome sequencing efforts, including the Earth Biogenome Project^7^ or the European Reference Genome Atlas^8,9^, generate vast genomic datasets, scalable and accurate functional annotation approaches are urgently needed.

Incorporating artificial intelligence (AI) techniques has enhanced the performance of prediction methods^10^. However, there is still room for improvement, especially when applied to full proteomes. Further developments benchmarked in CAFA datasets^10^, ranging from convolutional neural networks (CNN) deep learning models^11^ to protein language model-based approaches^12,13^, offer a promising alternative for decoding the ‘hidden biology’ within the dark proteome. We previously benchmarked those methods on model organisms and showed that protein language models (pLMs) predict more precise and informative GO terms than CNN-based methods, and have a higher coverage -proportion of annotated proteins in the proteome-than traditional homology-based methods, making them suitable for large-scale annotation^4^. Available tools that leverage pLMs are either hard to escalate to complete proteomes, install or customize for benchmarking or testing different models.

Here, we introduce FANTASIA (*Functional ANnoTAtion based on embedding space SImilArity*), a scalable pipeline that leverages pre-trained PLMs and similarity searches for large-scale functional annotation (Fig. 1). FANTASIA is a reimplementation of the GOPredSim algorithm^13^ focused on scalability for its genome-wide implementation and stability from scratch, together with the necessary steps for data preprocessing. By integrating an efficient implementation, parallel computing techniques, and a streamlined workflow, it significantly improves user-friendliness while maintaining prediction accuracy. First, given an input proteome or query proteins, it computes protein embeddings using state-of-the-art language models, like ProtT5^12^ or ESM2^14^. The list of proteins can be previously filtered by length, or by sequence similarity, to remove identical sequences within the input or close sequences in the references that can influence results during benchmarking. Then, it employs a robust embedding similarity-based approach to infer GO terms, accessing a precomputed database of embeddings with functional annotations. Next, it produces functional predictions using the closest hits or distance-based filtering methods to reduce noise and ensure robustness. Additionally, FANTASIA provides a flexible command-line interface, enabling seamless integration into diverse bioinformatics pipelines. See Methods for more details.

**Figure 1.**
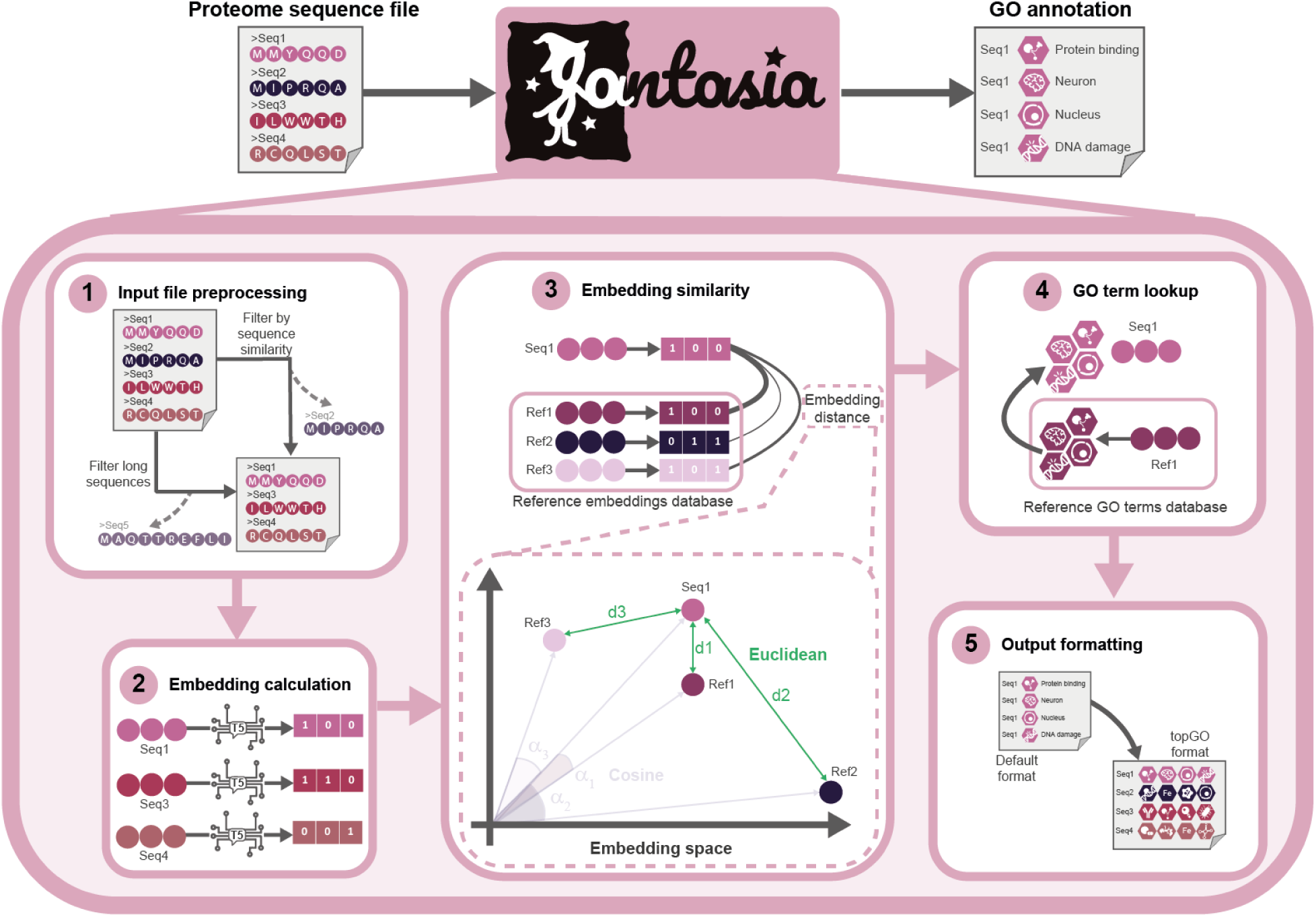
FANTASIA pipeline overview. Input proteomes are preprocessed to remove sequences based on length and sequence similarity. Then, embeddings are computed, and Euclidean embedding similarity is calculated against the reference database. Finally, it converts the standard GOPredSim output file to the input file format for topGO^15^ to facilitate its application in a wider biological workflow.

We previously benchmarked the original GOPredSim annotation algorithm in model organisms using the two provided pLMs (SeqVec and ProtT5) against two CNN-based models and a sequence profile-based method^4^. Our results showed pLMs are suitable for large-scale annotation and downstream analyses due to their increased annotation coverage (number of sequences annotated), term inference precision, and informativeness (or detail), while being able to recover accurate functional information from transcriptomic experiments.

Here, we expanded the analyses to 970 species (∼23 million genes) covering the full animal diversity. As these correspond to non-model organisms, there is no standard dataset available to do a proper benchmark as the previous one. Nevertheless, repeating some comparative analyses yielded the same results in terms of informativeness (S1) and annotation coverage, where traditional homology-based methods failed to annotate nearly half of the genes, particularly in less-studied phyla (Fig. 2a, S2). Moreover, we show that FANTASIA recovers less GO terms per gene (reducing the noise for downstream analyses, S3), recovers a moderately similar functional annotation to the homology-based one (S4), and exhibits minimal differences between using all isoforms (and combining all annotations) or just the longest (S5). In both methods, gene-level annotations remained highly similar, supporting the common practice of using the longest isoform in evolutionary studies while reducing computational costs.

**Figure 2.**
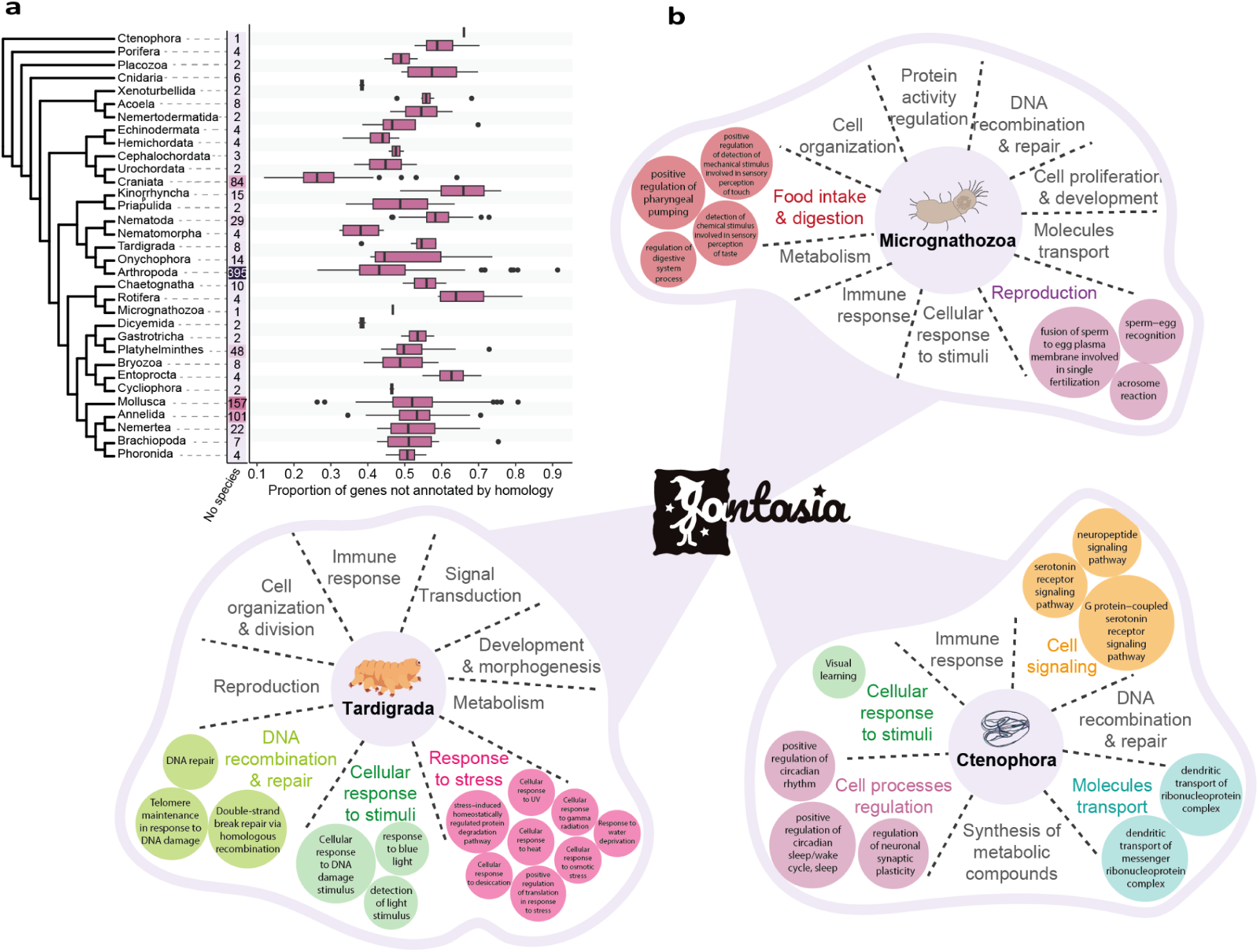
Functional annotation of the dark proteome across the Animal Tree of Life. **a,** Number of animal species per phylum included in this study (left) and the proportion of genes not annotated by homology per phylum (right). **b,** Enriched ‘hidden’ biological terms revealed by FANTASIA in three animal phyla. Large text summarizes key functions, while colored circles represent specific GO terms.

To show how FANTASIA can access previously ‘hidden’ biological information, we performed a per phylum GO enrichment analysis of the genes that were not annotated using homology-based methods. While 34.43% of GO terms were phylum-specific, 48 were common to all phyla (S6), mainly linked to viral response and immune function, both under adaptive evolution in animals. Newly annotated phylum-specific functions are not restricted to a specific group of functions but span diverse biological roles, including essential animal processes (S7). However, per-phylum GO enrichment analyses also revealed ‘hidden’ functions linked to unique biological traits. For instance, tardigrades showed stress-related functions that might help us understand their resilience; Micrognathozoa displayed functions tied to the pharynx and unexpected reproductive terms, hinting at undiscovered males or reproductive genes co-option; and ctenophores had newly annotated neuronal functions, shedding light on their unique nervous system (Fig. 2b, S8).

In the era of genomics, pLMs offer a promising complement to homology-based approaches for inferring gene functions in both model and non-model organisms. Here, we present FANTASIA as a reliable tool for inferring function in the coding region of newly-assembled genomes and demonstrate how these models can reveal previously ‘hidden’ functions across animals. After all, important biological functions can be found, even in the ‘darkest’ genes, if one has a way to turn on the light.

## Methods

### FANTASIA pipeline step-by-step

FANTASIA is a complete reimplementation of the GOPredSim algorithm^13^ for functionally annotating full proteomes based on GO term transference from protein embedding similarity. Building on the original method, our pipeline enhances usability by offering scalability to full proteomes, a more reliable installation process with minimized dependency conflicts, and an intuitive command line interface that simplifies parameter customization to facilitate usability. We provide two implementations of FANTASIA to accommodate different user needs. The first, FANTASIA v1, packaged within a Singularity container, was designed for scalability but presented challenges with dependency management. Based on user feedback regarding installation difficulties, we developed a second version (FANTASIA v2) that uses a vector database for improved stability and ease of deployment. The full list of differences between both versions is explained in the S9.

The FANTASIA pipeline consists of five steps: input preprocessing, protein embeddings computation, calculation of embedding similarity against an embedding reference database that contains functional annotations, GO terms transference from closest reference sequences embeddings, and output file formatting (Fig. 1).

#### Step 1: Input file preprocessing

FANTASIA allows the removal of long sequences and the application of a sequence similarity filtering using CD-HIT^16,17^. In FANTASIA v1, this preprocess is mandatory, ensuring that the pipeline can run smoothly by removing identical sequences in the input file and filtering out sequences longer than 5,000 amino acids. In contrast, FANTASIA v2 makes these steps optional and more flexible, allowing users to define maximum length and set a minimum sequence identity threshold (down to 50%). Unlike FANTASIA v1, the sequence similarity filter in FANTASIA v2 is conducted against the annotated reference database, and is only recommended for benchmarking purposes. If the latter filter is enabled, FANTASIA v2 concatenates the input dataset with the reference table before computing distances between embeddings (step 3 below). Then, for each query embedding, a comparison is performed against the entire reference table. During this process, embeddings corresponding to sequences that belong to the same cluster as the query are excluded, ensuring that no matches (hits) are made with proteins that exceed the specified sequence identity threshold. The use of these options depends on the intended application of the tool by the user, and are not recommended for full proteome functional annotation purposes. For details on parameter selection for different applications, refer to the pipeline documentation.

#### Step 2 : Embedding computation

FANTASIA computes the protein embeddings per model and per sequence. FANTASIA v1 only supports ProtT5^12^ (previously benchmarked in ^4^), while FANTASIA v2 also supports ESM2^14^ and ProstT5^18^. Embeddings are stored in HDF5 format for further use.

#### Step 3: Embedding similarity

FANTASIA then computes the distance between each input sequence embedding (query) and the embeddings in the reference database. The reference database in FANTASIA contains, for each reference protein, its pre-computed embeddings for the supported pLMs, and the GO term annotations that will be transferred in step 4 (see below). In FANTASIA v2, this information is stored as a reference vector database^19^ managed with PostgreSQL which also contains, for each reference protein, its sequence and metadata.

By default, Euclidean distance (*d*_*e*_) between embeddings *n* and *m* for a model with an embedding of embedding size *s* is computed with the following formula:

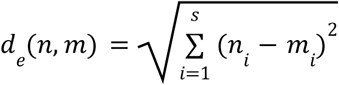

Where *s* represents the number of dimensions in the embedding space, which varies depending on the selected pLM: *s* = 1024 for ProtT5 and ProstT5, and *s* = 320 for ESM2.

Alternatively, FANTASIA v2 allows the selection of cosine similarity (*d*_*c*_), which is calculated using the formula:

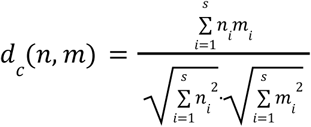

In FANTASIA v2, this step is done using pgvector, a PostgreSQL extension optimized for efficient similarity searches in high-dimensional embedding spaces^20^. By default, pgvector performs exact nearest neighbor search, ensuring perfect recall. However, it also supports approximate nearest neighbor search, which trades some recall for increased speed. Unlike typical indexes, adding an approximate index may yield slightly different query results. In our implementation, we use exact search to maximize accuracy, but we leave the option open for approximate search if faster retrieval is needed in future optimizations.

#### Step 4: GO transference

FANTASIA then transfers GO terms from the *k* closest embedding-bearing proteins in the database. By default, only the closest hit (*k* = 1) is used. In FANTASIA v2, users can define a distance threshold for each model that determines the maximum allowed distance between query and reference embeddings, which requires user optimization. FANTASIA v1 uses GO terms from the March 22^nd^, 2022 release (GOA2022), while FANTASIA v2 uses GO terms from the November 3^rd^, 2024 release (GOA2024).

#### Step 5: Output description and formatting

The standard output of FANTASIA gives the user information on the GO terms assigned to each input protein, along with the associated realiability index (see below). In FANTASIA v1, the output is a tab-separated file where each row represents a GO-input protein pair, with columns for the sequence accession, the GO term identifier, and the reliability index. In FANTASIA v2, the output is a comma-separated file (CSV) with 10 columns: 1) sequence accession (header); 2) GO term identifier; 3) GO category (F: molecular function, P: biological process, C: cellular component); 4) GOA evidence code for the reference annotation; 5) GO term description; 6) embedding distance (Euclidean or cosine) between the query and the reference; 7) protein language model used for the embedding generation; 8) UniProt ID of the reference protein bearing the closest embedding; 9) organism the target reference protein belongs to; 10) reliability index. By default, FANTASIA converts this standard output to the input file format for topGO’s GO enrichment^16^ to facilitate its application in a wider biological workflow in functional genomics. This feature can be disabled by the user in FANTASIA v2.

The reliability index (RI) was previously established in^13^, and is a transformation of the distance between the query *q* and hit in the database (*q*,*n*_*i*_) into a similarity scale, making it easier to compare between functional annotation results. Supported RI formulations:

For Euclidean distance (*d*_*e*_), RI is defined using the inverse similarity transformation as:

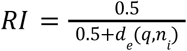

Where lower Euclidean distances yield higher confidence scores.

For cosine similarity (*d*_*c*_), RI is computed with the direct similarity measure:

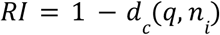

While both formulations produce values ranging from 0 to 1, they are not directly comparable, as they capture confidence in different ways. Users should exercise caution when interpreting RI scores across different similarity metrics.Additionally, the Euclidean distance is not inherently comparable across different pLMs, as it depends on the magnitude of the embedding vectors generated by each model. In contrast, the cosine similarity metric is more suitable for cross-model comparisons, as it primarily captures the relative orientation of embeddings rather than their absolute magnitude. Importantly, RI values do not necessarily correlate with the accuracy of the prediction, since relevant biological functions can still be recovered at scores within the second quantile distribution of RIs^4^.

FANTASIA can be executed on both GPUs or CPUs, depending on the available hardware, though GPU-based execution is recommended for improved speed (Fig. S9b). The computational complexity of the pipeline is O(n), where n is the number of sequences in the input dataset. Nevertheless, both versions differ on their speed, with FANTASIA v1 being seven times faster than FANTASIA v2. In addition, RAM usage also differs, with FANTASIA v1’s RAM usage increasing linearly with proteome size, while FANTASIA v2’s RAM depends on the reference database size. See S9 for more details on the comparison.

### Performance of FANTASIA (ProtT5) in proteomes across the animal tree of life

#### Input datasets and processing

We obtained the proteome files for 961 animals and 9 outgroup species from MATEdb2^21^ (File S1). This dataset comprises the longest peptide sequence (or isoform) per gene, totaling 23,184,398 sequences, derived from high-quality genomes and transcriptomes representing nearly all animal phyla. To assess the impact of annotating all isoforms versus only the longest, we created a subset of 93 species (File S1) to maximize representation across the Animal Tree of Life.

We annotated the functions (assigned GO terms) of the longest and all isoforms datasets using (1) FANTASIA v1 with the ProtT5 model and (2) eggNOG-mapper as a proxy for annotation based on homology. We used eggNOG-mapper 2.1.6^2^ as described in MATEdb2^21^. To ensure comparability, we used the datasets filtered by length and by removal of identical sequences, as provided by FANTASIA v1, as input for eggNOG-mapper. Homology-based results were then filtered to remove the GO terms that have a depth in the GO graph (DAG) lower than 2 (i.e. the terms are very general, close to or at the root of the DAG, and not informative). The difference in annotation coverage between the filtered and unfiltered homology-based annotations is negligible (Fig. S2, light purple).

When exploring the effect of using all isoforms for functional annotation, we first annotated all isoform sequences and then obtained per-gene annotations by collapsing the annotation of each isoform to retrieve a list of unique GOs.

In addition, we performed an extra RI-based filtering for FANTASIA results (see above for an explanation on RI). Thresholds for filtering were obtained from the source GOPredSim publication^13^ based on evaluations using CAFA3^10^: 0.35 for biological process, 0.28 for molecular function, and 0.29 for cellular component. It should be noted that these values were estimated based on the CAFA3^10^ F1 max comparisons datasets which do not necessarily scale to longer and more diverse datasets.

#### Information Content

To analyse GO terms according to depth within the GO graph, we estimated their Information Content (IC) using a custom script (https://github.com/cbbio/Compute_IC) which is defined as^22^:

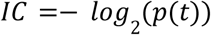

Where p(t), the probability of a term occurring in a dataset, is calculated as:

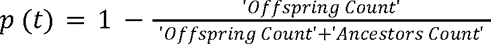

Offspring and ancestors count refer to the number of proteins that are assigned to that term given a reference Gene Product Association Data file (GAPD). In this case, the reference GAPD, or universe of proteins, combined all GAPD files for all species available.

Higher IC implies a more specific GO term while low IC values means that the GO term is more general.

#### Semantic similarity

To assess how similar two different annotations are, we obtained the lists of genes that were commonly annotated by both methods for the representative subset of species representing all phyla (n=93) and calculated for each gene their semantic similarity between the GO terms inferred independently for each of the 3 GO term categories. To be more conservative in our analysis, we used the most filtered dataset for each method (i.e., filtering by reliability index). We used pygosemsim (https://github.com/mojaie/pygosemsim) to estimate the Wang semantic similarity^23^ between protein-coding gene annotations using the Best-Match Average (BMA) method for gene products annotated with multiple GO terms. Semantic similarity (*sim*_*BMA*_) between 2 gene products (*g*1 and *g*2) is calculated as follows:

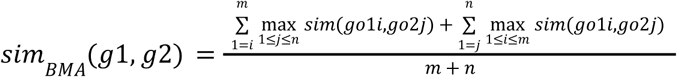

where *m* and *n* are the number of GO terms annotated in *g*1 and *g*2, respectively; and *go*1*i* and *go*2*j* are the GO term in position *i* (from 1 to *m*) from the *g*1 and the GO term in position *j* (from 1 to *n*) from the *g*2, respectively. *sim*_*BMA*_ Semantic similarity values range from 0 to 1, with 0 meaning no similarity and 1 meaning (almost) identical.

The Wang similarity method, unlike other metrics, is based on the GO DAG topology, making it more stable than others based on information content that rely on the available annotations.

#### GO enrichment for genes not annotated by homology-based methods

We obtained the list of genes that were not annotated by homology for each species and performed a GO enrichment analysis using topGO^15^ against the whole set of proteins of that species to identify which functions were being missed by homology. We used the GO terms predicted using FANTASIA. TopGO, in contrast to other GO enrichment methods, is based on the GO DAG, allowing us to take relationships between GO terms into account.

From the GO enrichment per species, we obtained a list of GO terms enriched per phylum by combining the results per species. For that, in the case of GO terms that were enriched in at least one species, we combined the p-values using the ordmeta method^24^.

We then identified the list of GO terms that were commonly enriched in all animal phyla. Among all common GO terms (Table S1), we focused on toxin activity (GO:0090729) to assess the performance of the functional annotation. Among all animals with venom, we selected the scorpion *Centruroides sculpturatus* to look into the genes with this GO term that were not annotated by homology. We used ToxinPred2^25^ to further filter this list and identified CSCU1_DN0_c0_g23215_i1.p1 (XP_023242874.1, uncharacterized protein) as a putative homolog of tachylectin-5 (XP_023224331.1). We further assessed this by predicting their structure with ColabFold^26^ and comparing them with the experimentally-determined structure of the horseshoe crab *Tachypleus tridentatus* tachylectin-5A (PDB 1JC9) using the PDB pairwise structure alignment tool. These results can be found in S6.

#### Data visualization

To visualize the significantly enriched GO terms per phylum and GO category, we first calculated the Wang semantic similarity matrix for each dataset and compared several clustering algorithms using the simplifyEnrichment R package^27^, and proceeded with the K-means algorithm (File S3). Then, to summarize the function of each cluster, we obtained their representative GO term by selecting the GO term with the lowest closeness value (the ‘farthest’) in the subset semantic similarity network for that cluster. Results were plotted in a TreeMap. We then combined the results of all phyla by assigning each significant GO term to the GO slim AGR (Alliance of Genomics Resources) subset using the map-to-slim.py script available in GOAtools^28^. Between 10 and 60% of GO terms per phylum were not assigned to a category and were further clustered as explained above. The GO representative terms of each of the 49 clusters were later manually assigned to the GO slim AGR classification and used to include unassigned GOs in the GO-GO category list. Final results were normalized per phylum (row) and visualized in a heatmap.

#### GO version

GO knowledgebase is regularly updated with new releases that remove obsolete GO terms or add new ones, which hinders comparison between analyses that use different versions^29^. Hence, we used the same version for all of the analyses whenever possible to improve comparability. For FANTASIA v1 animal proteome annotation, which is dependent on the GO release, we used the release of March 22^nd^ 2022 (GOA2022) and was compatible with the previous benchmark. This same release was used for the GOslim subsets. For information content and semantic similarity analyses, we used the release of June 11^th^ 2023.

### Computational resources

FANTASIA was tested in several computing environments. The performance of FANTASIA v1 was assessed in the Finisterrae III HPC (FT3, Centro de Supercomputación de Galicia, CESGA) using CPU nodes, and NVIDIA A100 GPU nodes with 32 CPUs per node and a maximum of 250 GB RAM, CUDA version 11.0. FANTASIA v2 was evaluated on (1) a desktop workstation at CBBIO lab wtih an AMD Ryzen 9 5950X 16-core processor CPU (128 GB RAM), and a NVIDIA GeForce RTX 3090 Ti GPU (24 GB VRAM), CUDA version: 12.4; and (2) an HPC with NVIDIA A100-SXM4-80GB with CUDA 12.1, utilizing a single GPU, 50 CPU cores, and 100 GB of RAM. Animal proteomes GO annotations were performed in DragoHPC cluster (Spanish National Research Council, CSIC) using nodes with an NVIDIA Ampere A100 graphic card with 512 GB and Dual Intel Xeon Gold 6248R 24C 3.0GHz processors. Only one species (*Panulirus ornatus*) was annotated using CPUs in FT3 due to the limitation in the maximum GPU nodes RAM memory.

## Code availability

Both versions are functional and available. FANTASIA v1 pipeline used for the supplementary analyses is included in a Singularity (v3.8.3+231-g9dceb4240) container available on GitHub (https://github.com/MetazoaPhylogenomicsLab/FANTASIA). FANTASIA v2 pipeline is available on GitHub (https://github.com/CBBIO/FANTASIA). Steps followed for the creation of several intermediate files not available on Zenodo, as well as additional scripts used for the analyses and creating figures are available on GitHub (https://github.com/MetazoaPhylogenomicsLab/Martinez_Redondo_et_al_2024_Dark_Proteome_Animal_Tree_Of_Life).

## Data availability

All proteomes, homology-based annotations and FANTASIA.V1 (ProtT5 per-protein embeddings and annotations) are available in MATEdb2^21^. Information content, semantic similarity and other results are provided in Zenodo^30–35^. The complete list of the Zenodo files, as well as the supplementary files S1-4 can be found on GitHub (https://github.com/MetazoaPhylogenomicsLab/Martinez_Redondo_et_al_2024_Dark_Proteome_Animal_Tree_Of_Life).

## Acknowledgments

GIMR, AMR, and RF acknowledge funding and support from the LifeHUB/CSIC research network (PIE-202120E047). AMR and RF acknowledge funding from the OSCARS project, which has received funding from the European Commission’s Horizon Europe Research and Innovation programme under grant agreement No. 101129751. GIMR acknowledges the support of Secretaria d’Universitats i Recerca del Departament d’Empresa i Coneixement de la Generalitat de Catalunya and ESF Investing in your future (grant 2021 FI_B 00476). RF acknowledges support from the following sources of funding: Ramón y Cajal fellowship (grant agreement no. RYC2017-22492 funded by MCIN/AEI /10.13039/501100011033 and ESF ‘Investing in your future’), the Agencia Estatal de Investigación (project PID2019-108824GA-I00 funded by MCIN/AEI/10.13039/501100011033), the European Research Council (this project has received funding from the European Research Council (ERC) under the European’s Union’s Horizon 2020 research and innovation programme (grant agreement no. 948281)), the Human Frontier Science Program (grant no. RGY0056/2022) and the Secretaria d’Universitats i Recerca del Departament d’Economia i Coneixement de la Generalitat de Catalunya (AGAUR 2021-SGR00420). AMR, FPC and IB acknowledge support from MDM-2016-0687 and PID2021-127503OB-I00 funded by MCIN. We also thank Centro de Supercomputación de Galicia and CSIC for access to computer resources (CESGA and DRAGO respectively).

## Author contributions

GIMR, AMR and RF designed the study. GIMR performed the main analyses, created the FANTASIA v1 singularity container, and designed the figures. GIMR and IB wrote the original FANTASIA code and the pipeline to automate the methods. FMPC developed the FANTASIA v2 implementation. IC, and JMF optimized development, documentation, and extensively tested the tool and conducted statistics. MVV assisted with data analysis. GIMR, AMR and RF wrote the first version of the manuscript. RF and AMR provided resources, the rationale and concepts behind the work, and supervised the study. All authors revised and approved the final version of the manuscript.

## Supplementary information

### S1. Comparison of the level of detail of GO annotations between methods

To assess the level of detail of the non-homology-based annotations, we compared the information content (IC) of their GO terms (see Methods). Our results showed that FANTASIA and homology-based annotations are specific and therefore detailed in all categories, with FANTASIAm values being slightly higher (Fig. S1a). IC did not differ among phyla (Fig. S1b).

**Figure S1:**
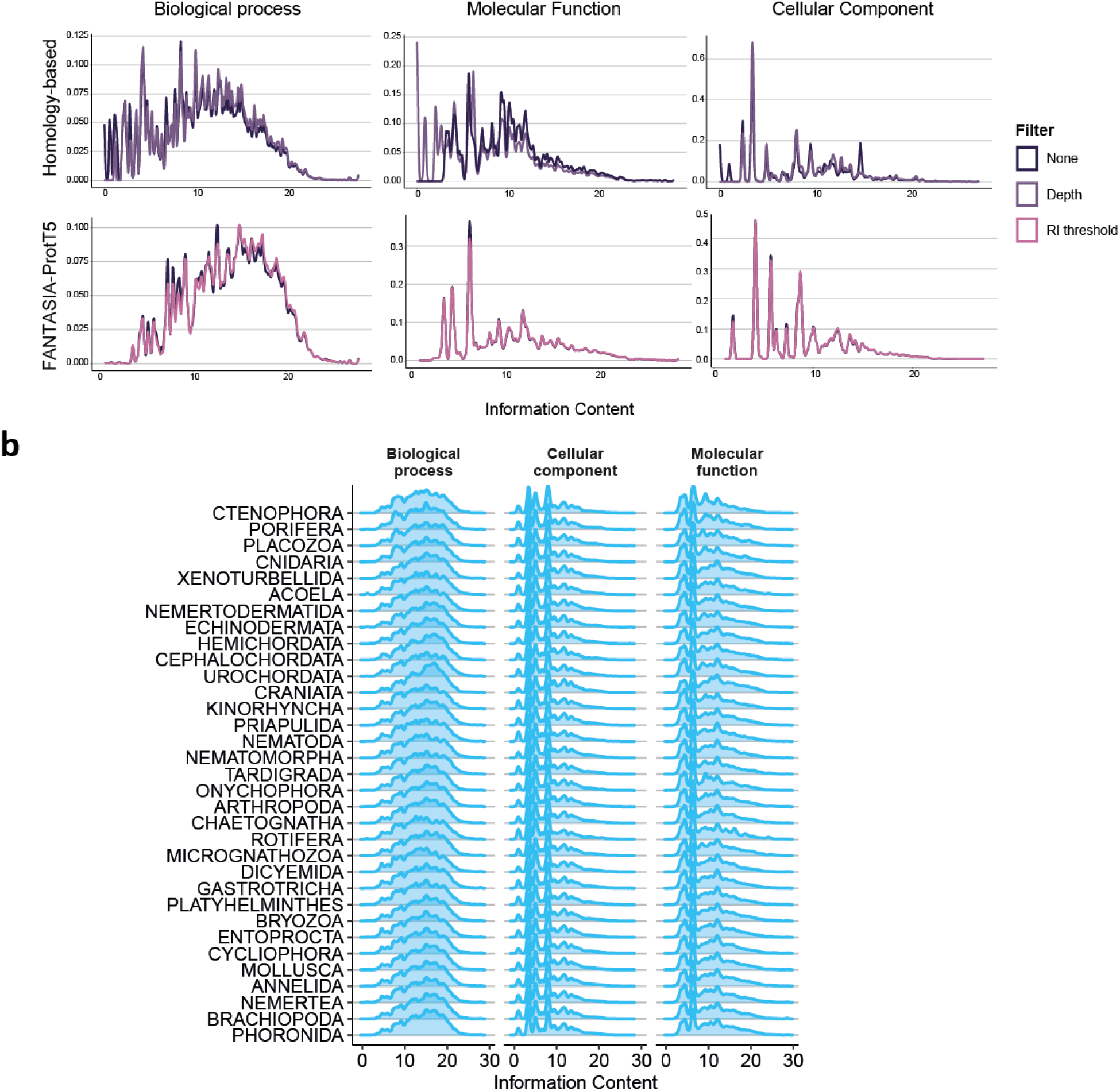
Comparison of information content (IC). **a,** Comparison of the IC of the annotations made by the different methods, for every filtering and GO category. **b,** IC per phylum for all 3 GO categories inferred by FANTASIA for the reliability index filtered dataset.

### S2. Effect of filtering by reliability index on annotation coverage per method

In total, after this filter, the number of genes annotated by FANTASIA-ProtT5 was reduced, losing 11.32% (Fig. S2a, pink). Per-phylum proportion of annotated genes after filtering followed the same pattern as in homology-based annotation, with better-annotated phyla having a higher proportion (Fig. S2b). Thus, in terms of the proportion of annotated genes, FANTASIA-ProtT5 annotates the highest number, even when applying a restrictive filter, outperforming homology-based methods.

**Figure S2:**
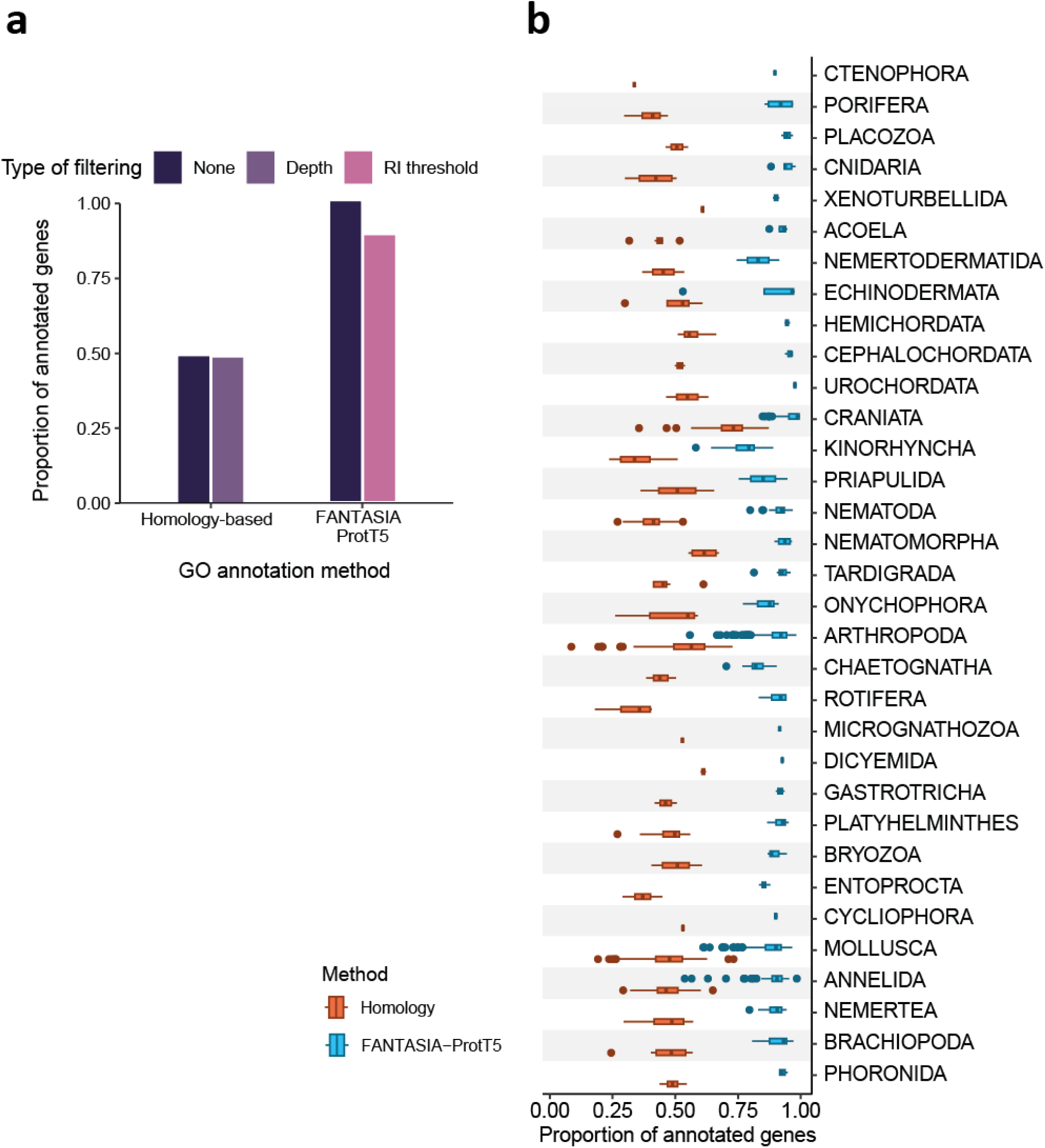
Comparison of the proportion of annotated genes with homology, and FANTASIA-ProtT5 using different filters. **a,** Proportion of genes annotated without applying any filter (dark purple), filtering by depth (light purple), and filtering by reliability index (RI, pink)**. b,** Proportion of annotated genes per animal phylum with the most filtered datasets.

### S3. Comparison of the number of GO terms per protein between methods

Regarding the number of GO terms assigned per gene, eggNOG-mapper assigned overall higher numbers of GO terms, with a median of 74 and reaching maximum values higher than 1200 for the biological process. Conversely, the median number of GO terms assigned to each gene by FANTASIA was 2, with the maximum being lower than 200 for the biological process. In both methods, the number of biological process GO terms was the highest of the three categories, followed by the molecular function and the cellular component, in this order. These patterns were constant regardless of the filtering method used, even though filtering by depth resulted in a slight reduction of the maximum and mean values (Fig. S3).

**Figure S3:**
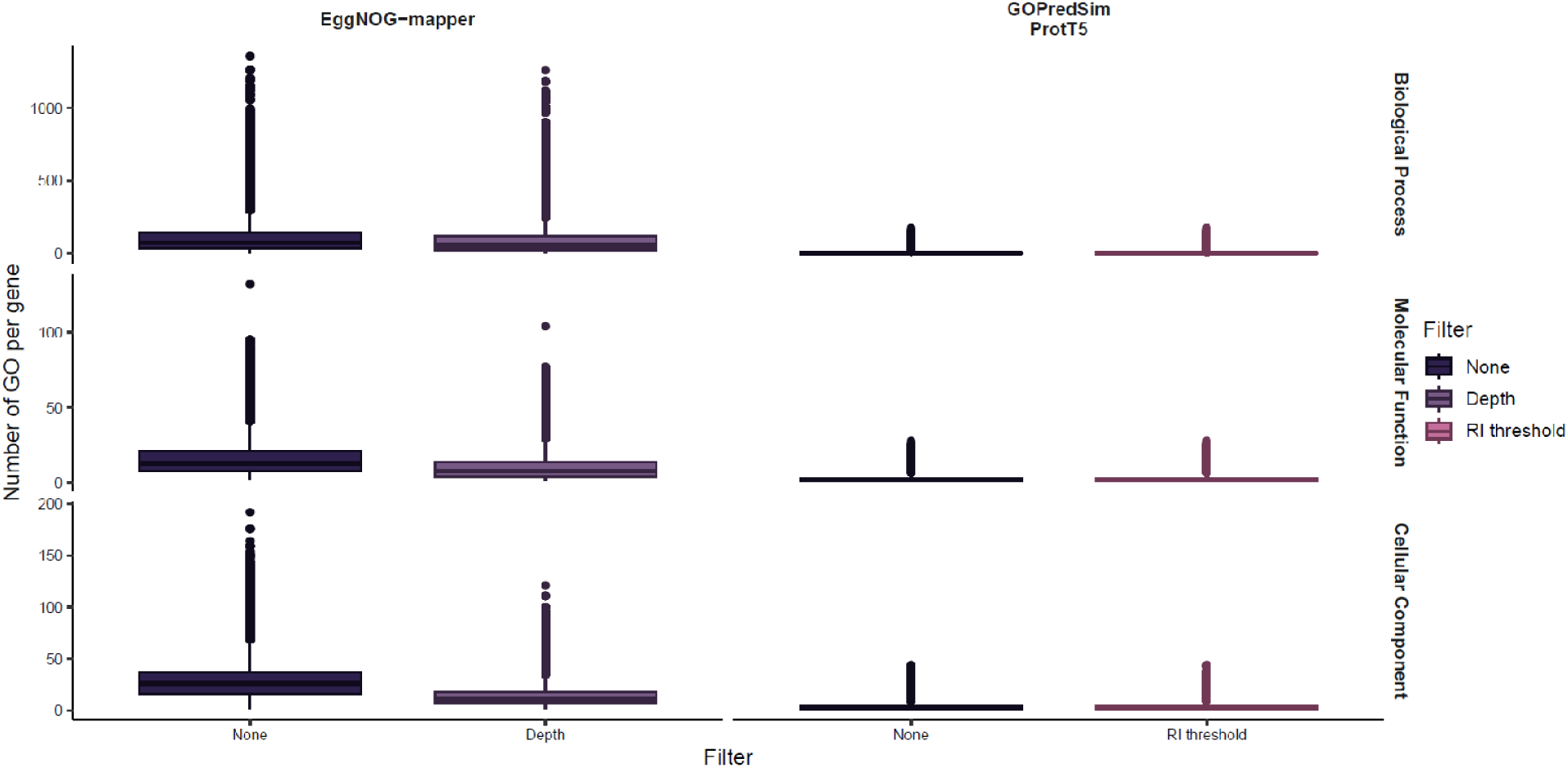
Number of GO terms of each GO category per gene in all tested functional annotation methods.

### S4. Semantic similarity of the GO annotations between methods

We assessed how similar the annotations of each of the methods were against the homology-based annotations. Briefly, the semantic similarity between two sets of GO terms (gene annotations) was estimated by aggregating the individual GO terms Wang semantic similarity (which considers how close are two GO terms in the GO DAG) using the Best-Match Average strategy (see Methods). For this, we used only the subset of genes that were annotated with both methods. Our semantic similarity results showed that the annotations of FANTASIA (ProtT5)varied to some extent compared to those based on homology. Annotations obtained for biological process were less similar than the ones obtained for molecular function and cellular component for all comparisons (Fig. S4a), which is consistent with our previous results in model organisms^4^, and with the higher diversity of biological process GO terms^36^. The lower semantic similarity might be explained by the higher number of GO terms per gene in the homology-based annotation (denominator in the formula).

Moreover, semantic similarity did not differ among phyla (Fig. S4b).

**Figure S4:**
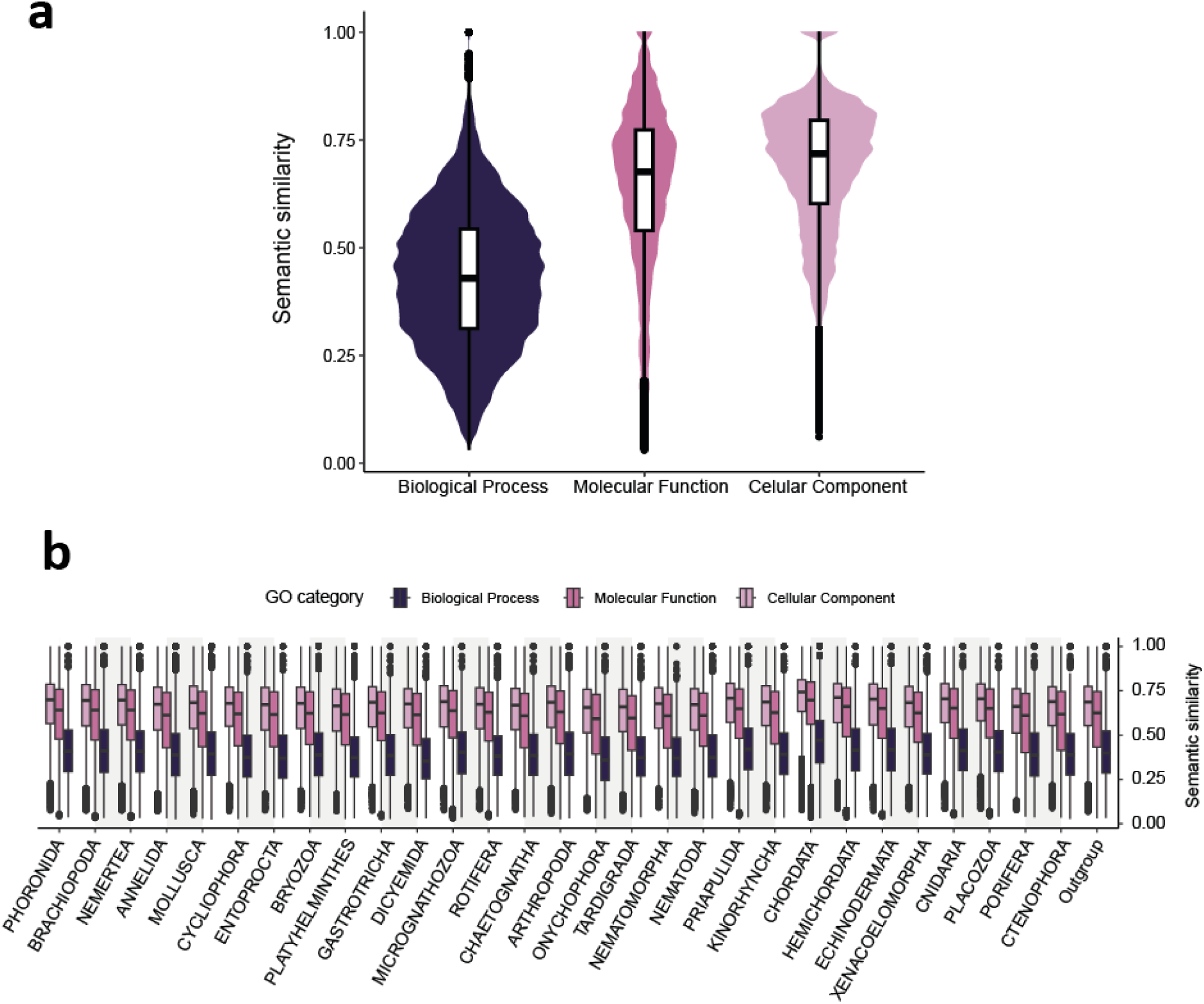
Semantic similarity between the gene annotations from FANTASIA-ProtT5 (filtered by RI) and homology-based annotations for all 3 GO categories. **a**, Full dataset comparison. **b**, Per phylum.

### S5. Assessing the impact of using all isoforms for per-gene functional annotation

In evolutionary studies of non-model organisms, it is common practice to use the longest isoform of each gene as a proxy for the whole gene, particularly in phylogenomic studies aiming at species tree reconstruction or gene repertoire evolution, to reduce the redundancy of the sequences included in the dataset and have only one sequence representing each gene. This is commonly preferred to the use of the consensus sequence per gene as these are artificial constructs that do not have a real biological function. Still, it remains unclear whether ignoring shorter isoforms affects the prediction of gene function or not. To see how this annotation may change, we annotated the functions of a subset of 93 proteomes representing all animal phyla including all isoforms per gene and compared it with the annotation previously inferred using only the longest isoform. Our results show that FANTASIA, with and without filtering by reliability index, has fewer differences in the numbers of annotated genes when either considering all isoforms or just the longest, than homology (Fig. S5a, secondand third panel, respectively).

We compared how similar the annotations were at the gene level when annotating only the longest isoform or all of them per gene. More than half of the genes had the same or very similar annotation for both methods (Fig. S5b). In addition, the patterns of information content were highly similar using either the longest (Fig. S1c) or all isoforms (Fig. S5) when comparing the different methods. Overall, these results indicate that the annotations at the level of gene are very similar if we annotate only the longest isoform per gene or all of them for virtually all the methods tested, which will reduce computational efforts on one side and allow the exploration of proteomes reported in the literature that include only the longest isoform per gene on the other, as is common in phylogenomic studies of non-model organisms.

**Figure S5:**
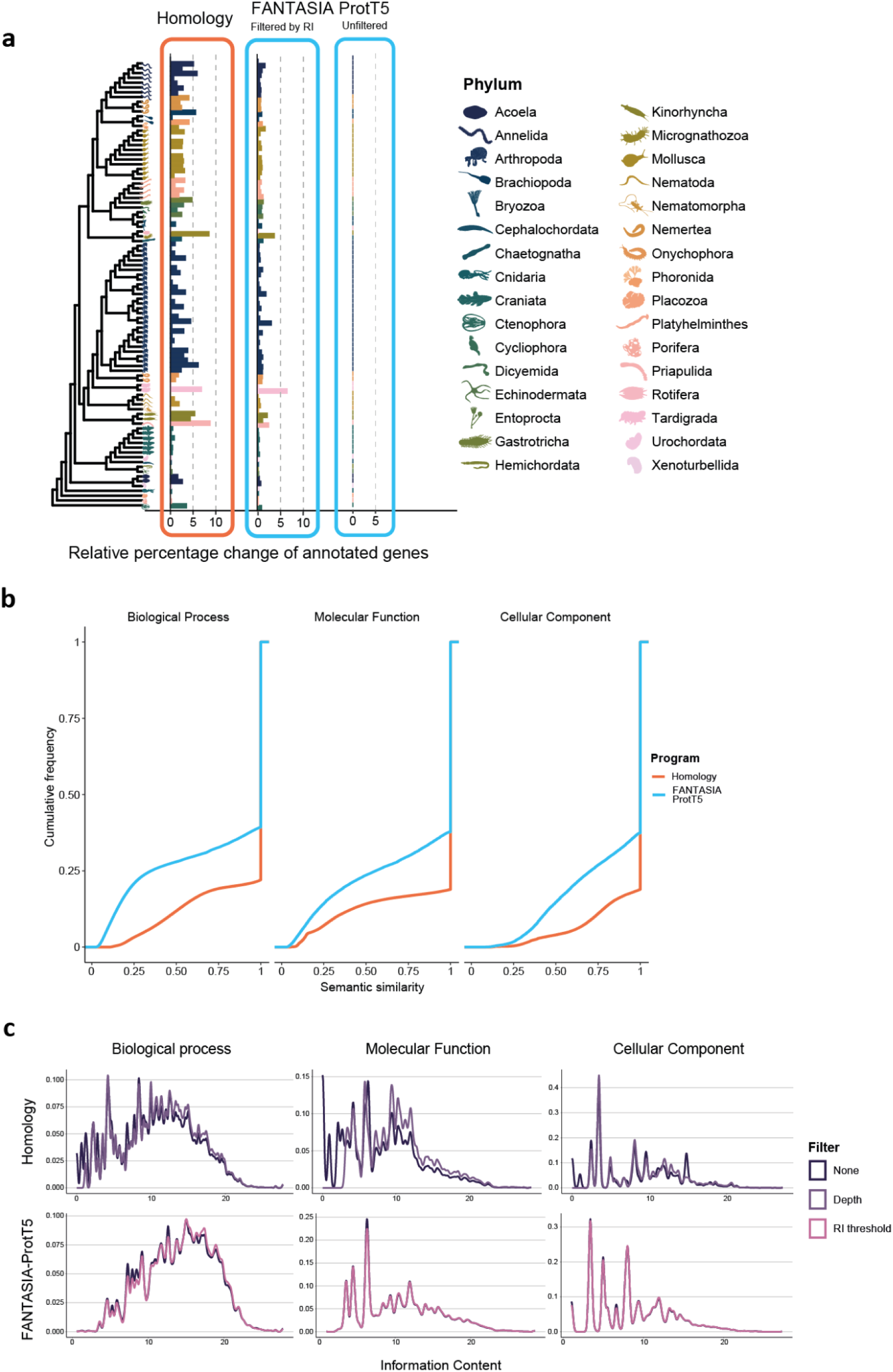
Per-gene functional annotation differences when using all isoforms or just the longest isoform. **a,** Percentage change of genes annotated when using all isoforms compared to genes annotated using the longest isoform. Results are shown for the most filtered datasets, i.e. FANTASIA-ProtT5 filtered by reliability index. **b,** Semantic similarity when comparing per-gene functional annotations obtained using all isoforms with the annotations obtained using the longest isoform**. c,** Comparison of the Information Content (IC) of the per-gene isoform-based annotations made by the different methods, for every filtering and GO category.

### S6. Common GO terms in the dark proteome across the Animal Tree of Life

To understand which functions have genes that are not annotated with the traditional homology-based methods, we performed a GO enrichment analysis per phylum of the functions inferred by FANTASIA for the genes not annotated by homology (see Methods). While 34.43% of all GO terms were found to be phylum-specific, 48 GO terms were common to all phyla (Table S1, Fig. S6a). Common GO terms were mainly related to viral response and the immune system, functions that have been proven to be under adaptive evolution in animals^37–39^.

For instance, one common GO term was GO:0090729 (toxin activity), revealing a set of candidate genes that may be relevant to understanding the defense or hunting mechanisms of animals, as toxins and resistance to them is another trait usually under strong selection^40^. As an example, we identified an undescribed tachylectin-5-like protein in a scorpion species annotated with this GO term (Fig. S6b, see Methods). Tachylectin-5 is a protein from the hemolymph of horseshoe crabs involved in its innate immune response against pathogens^41^, which makes their ‘blue blood’ indispensable for drug testing. Related proteins have been found to be expressed in the venom glands of some spiders^42^, suggesting a putative toxin-like role in scorpions.

**Table S1:**
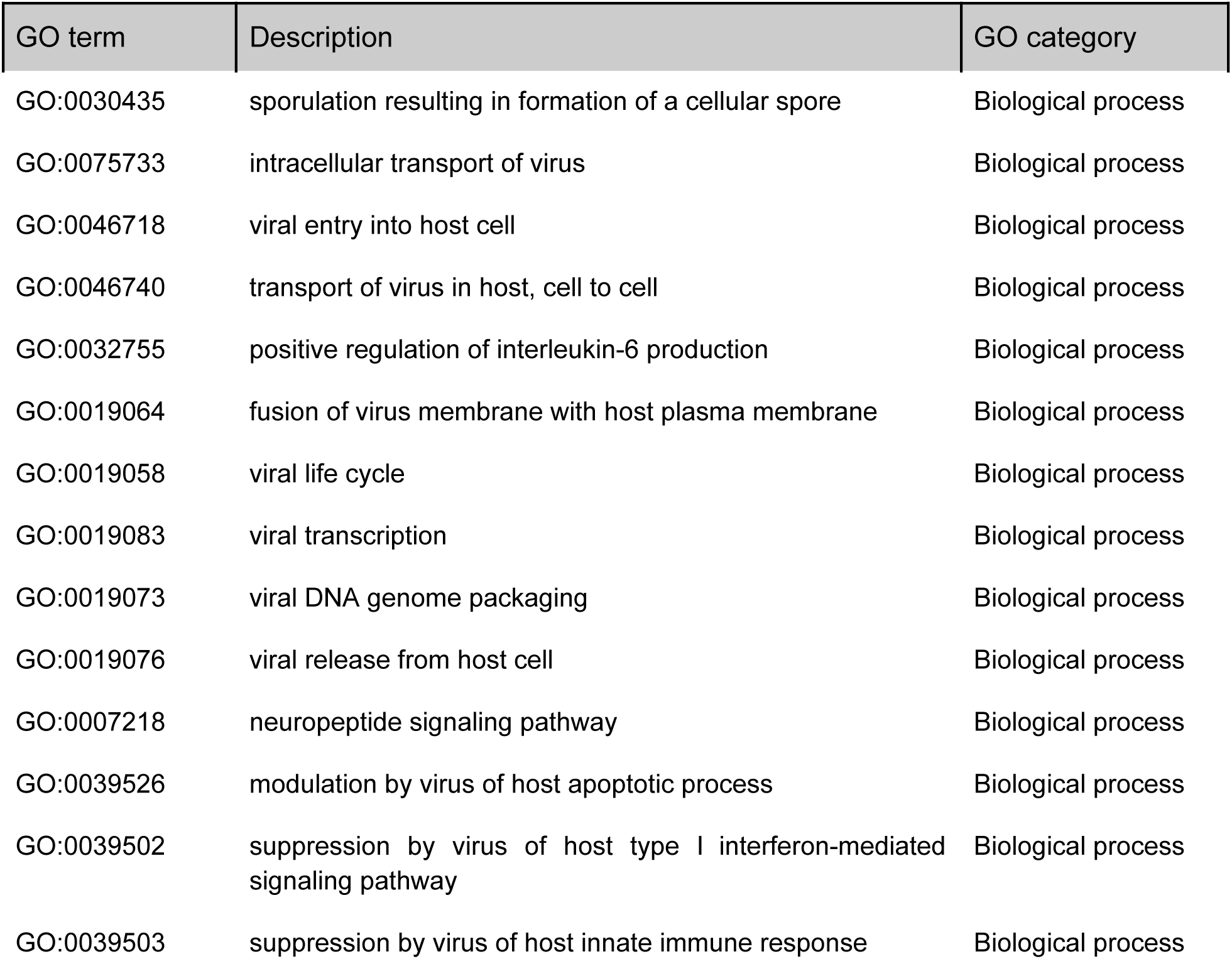
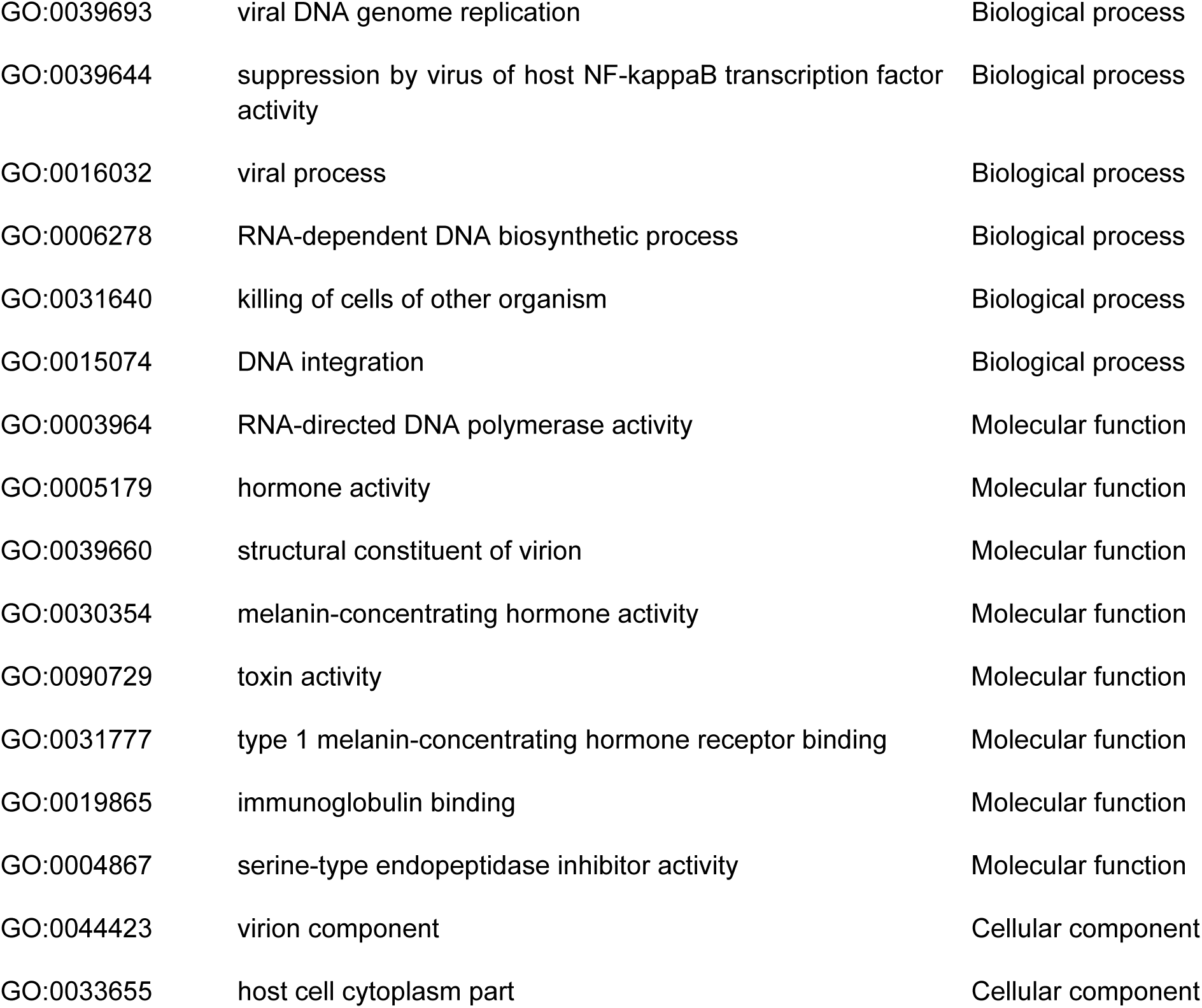
GO terms significantly enriched in the ‘dark proteome’ that were common to all animal phyla.

**Figure S6:**
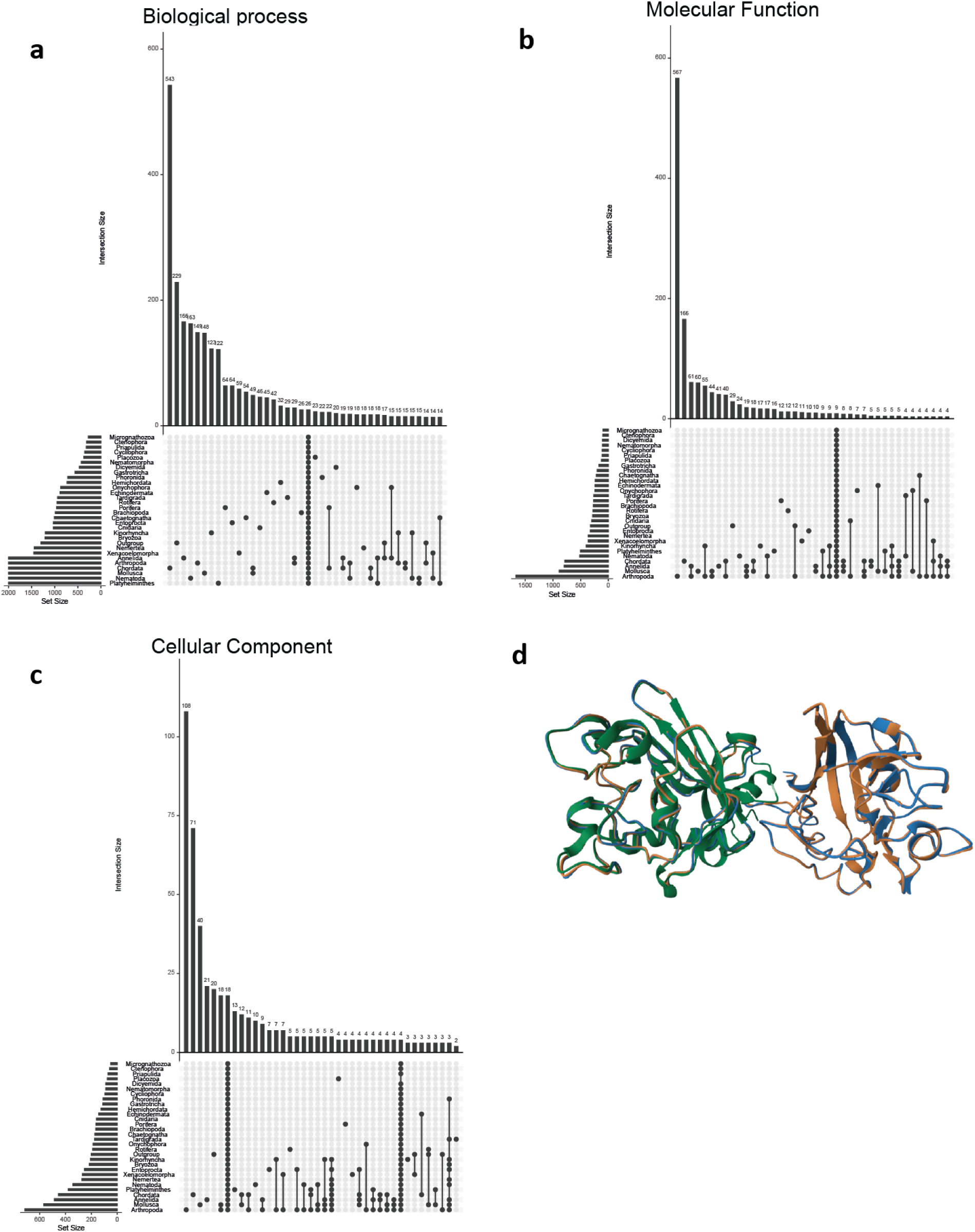
Common GO terms among all animal phyla. **a,** UpSet plot for biological process. **b,** UpSet plot for molecular function. **c,** UpSet plot for cellular component. **d,** Structure alignment of *C. sculpturatus* uncharacterized protein XP_023242874.1 (in orange), tachylectin-5A-like XP_023224331.1 (in blue), and *T. tridentatus* 1JC9 (in green). RMSD against the uncharacterized protein is 0.44 and 0.53. respectively.

### S7. Non-common GO terms in the dark proteome across the Animal Tree of Life

Among non-common GO terms, we found plant and fungi-specific functions (e.g. “Photosynthesis”). To see the extent of getting these putative “artifacts” in our results, we traced back in our subset of 93 animals how many GO terms were transferred from non-animal species in the lookup reference dataset. However, tracing back the taxonomic origin of the GO terms for our subset of 93 animals, we found that 78% out of the 64% matching proteins genes were of animal origin, being these non-animal GO terms a minority mostly affecting biological process terms. The remaining percentage not being straight-forward to assign, something we fixed in the new FANTASIA implementation. Regardless, these few cases could still be interesting (once ruled out any potential methodological artifact with further analyses) since they might point to convergent changes at the functional level across distant taxonomic scales, i.e., proteins that are similar (in their molecular function) but are involved in perform different processes; or deep homologs hard to identify as such with traditional methods. For instance, some GO terms from gene CG11373 in *D. melanogaster* (Table S2) were transferred from a fungal species, as deduced from some fungi-specific GO terms (GO:0140538 -negative regulation of conjugation with zygote-; and GO:0030435 -sporulation resulting in formation of a cellular spore-). This gene lacks GO term annotation in Flybase^43^, but the FANTASIA-predicted functions (Table S2) are consistent with its described expression in the male germline and testis somatic cells.

Next, we summarized the enriched functions by assigning each GO term to an Alliance for Genomics Resources (AGR) GO Slim GO for a simplified visualization of their function (Fig. S7a). We found that for all animal phyla, most of the enriched biological processes functions newly unveiled were related to response to stimuli, followed by immune system and developmental processes, signaling, and the establishment of localization (Fig. 7a, left). Molecular function and cellular component functions revealed that these newly annotated proteins have mainly catalytic activity or bind other molecules as parts of protein complexes (Fig. 7a, center, and right, respectively). This suggests that genes not annotated by homology are not restricted to a specific group of biological functions and might be previously unexplored proteins involved in widely studied essential animal processes.

**Table S2:**
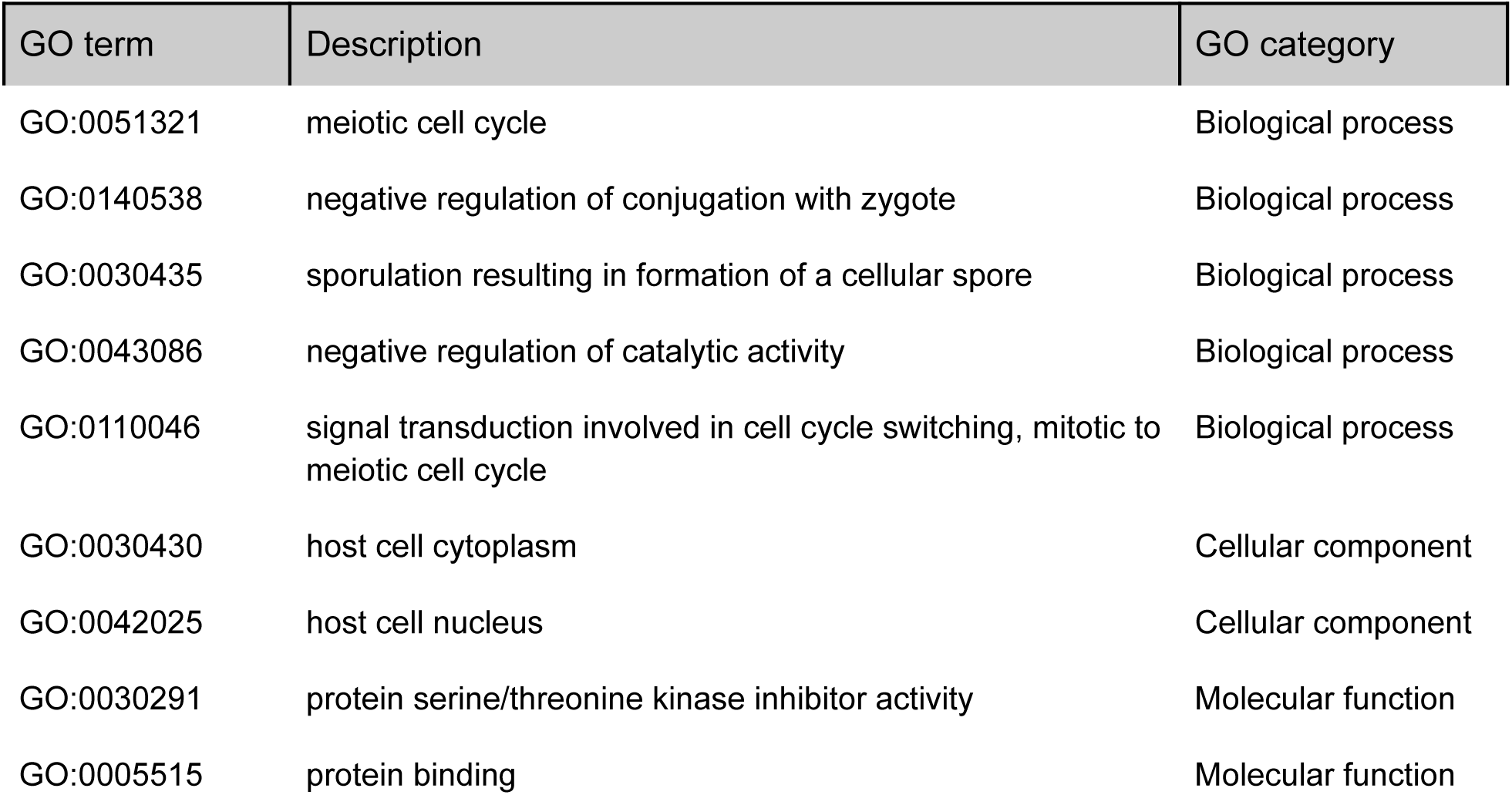
FANTASIA GO terms for gene CG11373 in *D. melanogaster*.

**Figure S7:**
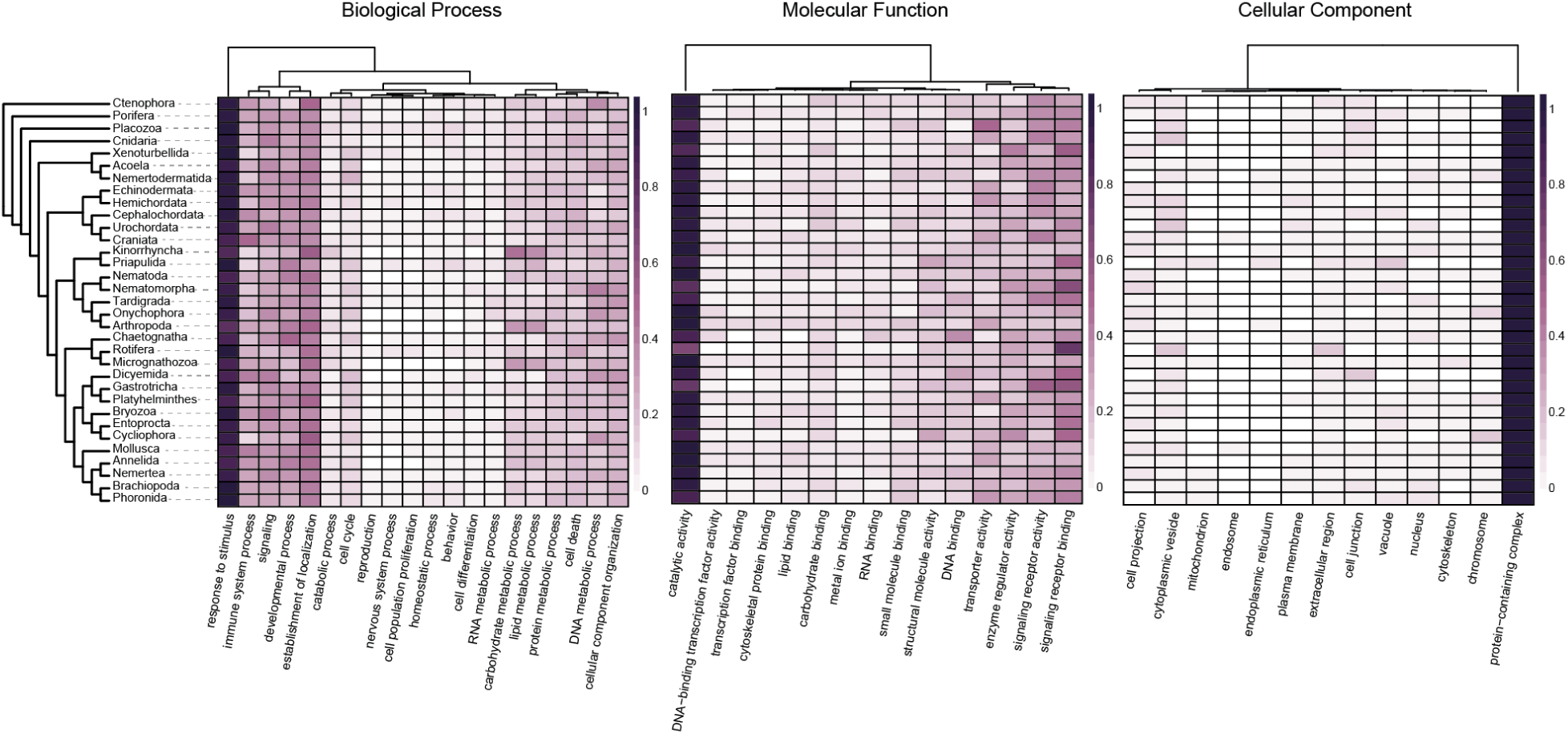
Summarized functional annotation of the dark proteome across the Animal Tree of Life. Per-phylum normalized number of enriched GO terms assigned to AGR GO slim terms.

### S8. FANTASIA per-phylum predicted GO terms reveal the ‘hidden biology’ in the dark proteome across the Animal Tree of Life

Phylum or lineage-specific enrichment results also show interesting hidden functions related to their unique biological characteristics. For instance, tardigrades, commonly named water bears or moss piglets, are known for their high resilience to environmental stressors, such as desiccation or high levels of radiation^5^. Tardigrades contain newly annotated genes related to the response to stressors such as UV-A, osmotic stress, or heat, which may help us to understand how these animals can survive in such extreme conditions (Fig. S8a). Another example is the enigmatic monotypic phylum Micrognathozoa (Fig. S8b). These microscopic animals are known for their complex pharyngeal apparatus^44^. Here, we recover functions enriched in pharyngeal pumping and digestive system regulation, together with touch and taste perception, which may help us to understand the origin of such a complex pharynx. In addition, no males have been ever described and micrognathozoans are assumed to be parthenogenetic^44^. Nevertheless, our results revealed GO terms related to sperm and egg fusion, suggesting that males may exist but have not been observed yet (probably due to their small size as occurs in rotifers), and thus they may have sexual reproduction. Another possibility, if we consider them asexual, is that the genes that are involved in these sexual functions in other animals have not been lost in this phylum and were co-opted for another mechanism that shares some similarities at the molecular level. The last example is the ctenophores (Fig. S8c), whose putative position in the Animal Tree of Life as the sister lineage to the rest of animals^45^ and their unique complex syncytial nervous system^46^ suggests that neurons may have evolved twice in animals. Here, newly inferred functions previously unannotated exposed GO terms related to neuronal activity (i.e., regulation of neuronal synaptic plasticity, neuropeptide signaling, dendritic transport of messenger ribonucleoprotein complex, and regulation of potassium ion transport), providing new putative annotated candidate genes that may shed light into the evolutionary origin of the nervous system in ctenophores. GO enrichments of the ‘hidden proteome’ of each animal phyla (for all three GO categories) are not discussed here due to space constraints, but we encourage the readers to inspect them in the File S2, as we believe they can serve as a very valuable source of candidate genes to further explore with other methodologies for hypothesis-driven functional testing.

**Figure S8:**
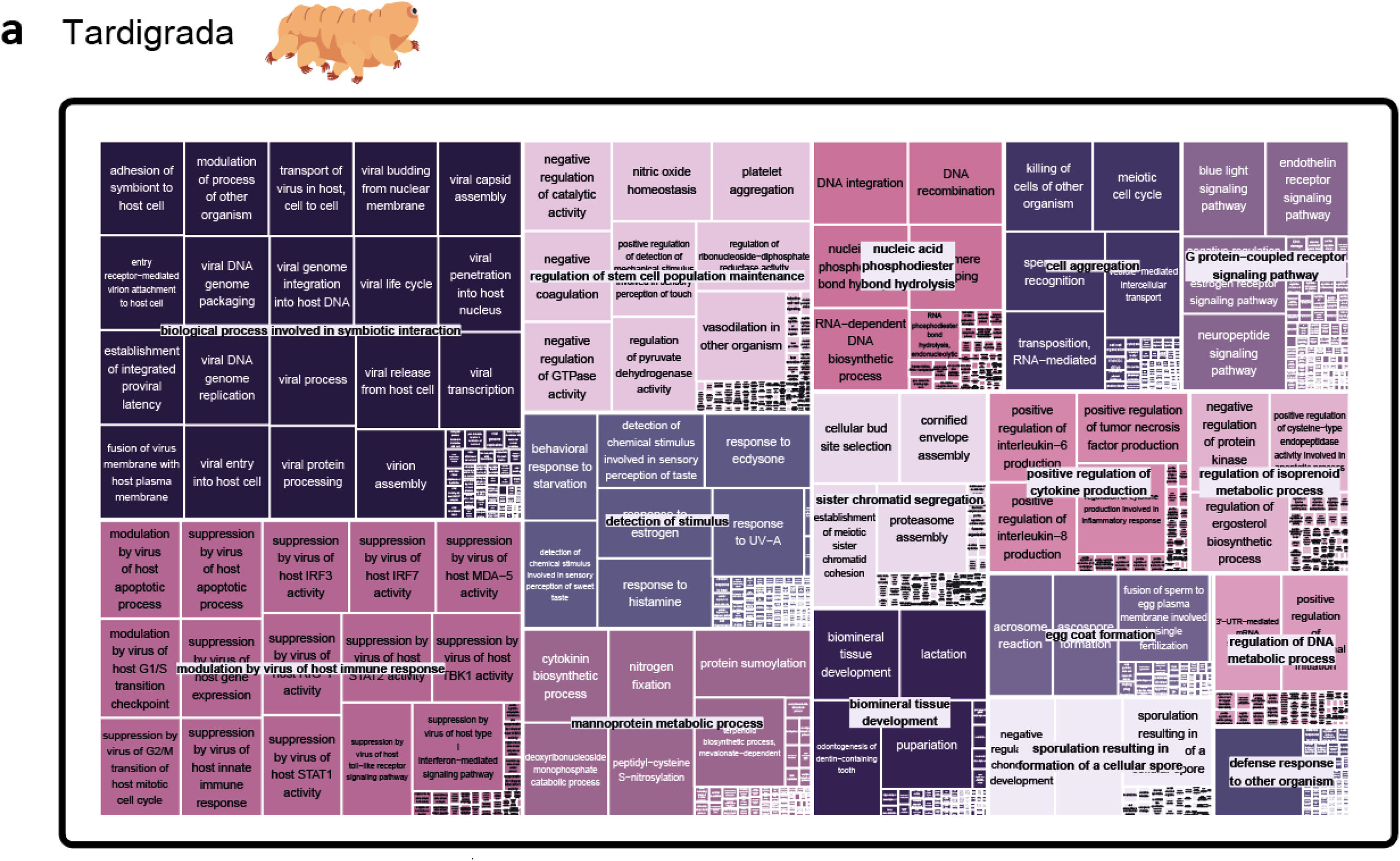
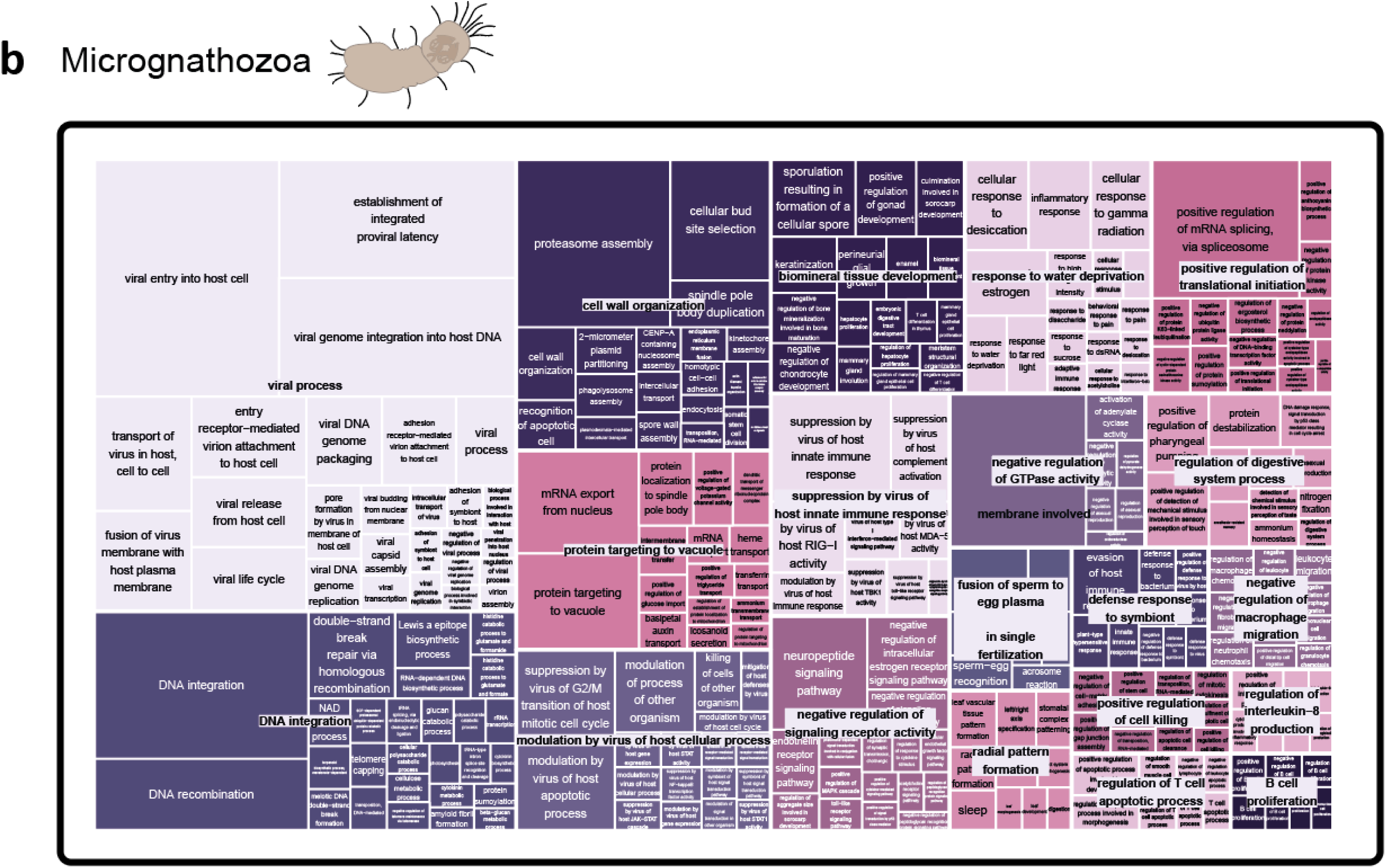
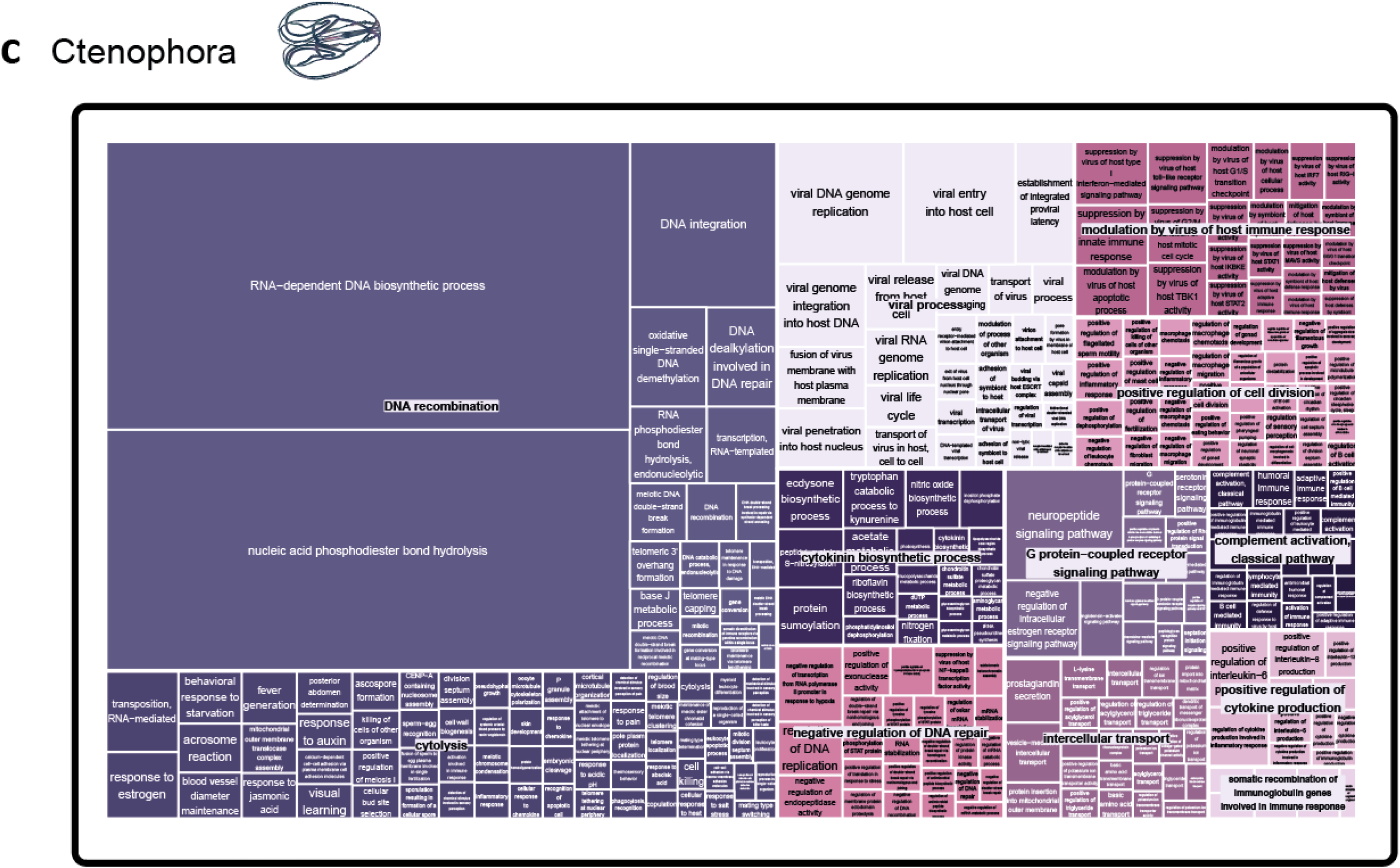
Treemaps showing enriched biological process GO terms in some animal phyla. The size of the square is proportional to the p-value of the enrichment. **a,** Tardigrada. **b,** Micrognathozoa. **c,** Ctenophora.

### S9. Differences between FANTASIA v1 and v2

While FANTASIA v1 (implemented in a Singularity container) has been already used by multiple users since it was created and made available on GitHub, it faced issues primarily related to dependency conflicts when installing on some computing systems. These reasons prompt us to create FANTASIA v2, which fixes those limitations. The core algorithm remains identical in both versions, with differences in the preprocessing of input files, output format, the flexibility of parameter selection, the method for similarity searches, the expansion of supported embedding models, updates to the the GO annotation database, embedding precision computation, and resources usage. More specifically:

1. In FANTASIA v1, sequences longer than 5000 amino acids were automatically removed. In FANTASIA v2, long sequences are supported and can be optionally removed by selecting a filtering threshold.
2. In FANTASIA v1, CD-HIT^16,17^ was always applied to remove proteins with 100% sequence identity within the input proteome, without an option to disable it. Keeping those sequences leads to execution errors. In FANTASIA v2, identical sequences can be safely retained. Morever, the sequence similarity filtering step is now optional and is conducted against the annotated reference database (allowing a minimum identity threshold of 50% between query and reference), and is only recommended for benchmarking purposes
3. FANTASIA v2 introduces extensive parameter customization, allowing users to define multiple aspects of the workflow, including the selection of embedding models, similarity search thresholds, and other benchmarking configurations. The full list of customizable parameters is available in the documentation.
4. FANTASIA v2 expands beyond the ProtT5 model used in FANTASIA v1, incorporating two additional state-of-the-art protein language models: ESM2, and ProstT5.
5. FANTASIA v1 used the GOA2022 release (March 22^th^, 2022) as a reference, while FANTASIA v2 uses GO terms from the November 3^rd^, 2024 release (GOA2024).
6. One of the main differences between versions is the numerical (floating-point) precision (FP16 and FP32 for FANTASIA v1 and v2, respectively). In FANTASIA v1, FP16 (‘half precision’ option in the configuration file) was selected following the recommendations from the *bio_embeddings*^47^ library authors, as they noticed a decrease in memory consumption while having a negligible effect in their benchmark results. To demonstrate that the algorithm provides coherent results regardless of its implementation and numerical precision, we tested the ProtT5 annotations and their derived RI values provided by both FANTASIA versions on 45,812 mouse proteins (File S4) with annotations across the three ontologies. No major inconsistencies were found in the RI values. Both distributions are very similar with most annotations at the high scores. In addition, the majority annotations exhibit small differences in the scoring by both versions (∼82% of BP, and ∼75% of CC and MF, exhibit differences < 15% of the relative score, for details S9). While these small differences may be due to how the distance is calculated and the floating-point precision, with FANTASIA v2 being more conservative, extreme differences are most likely to different coverage by different GOA versions where annotations change.
7. Computational resource usage differs between FANTASIA v1 and v2. In terms of execution time, FANTASIA v1 is generally faster than FANTASIA v2 for both embedding computation and GO transference. For ProtT5 embeddings generation on 87,492 mouse sequences (taxonomy id: 10090, time-stamped 2025-03-17), FANTASIA v2 maintains a fixed speed of 83.05 ms/protein on GPUs, whereas FANTASIA v1 shows greater variability when tested on a variety of proteomes of different size^30^ (∼14.71–120.48 ms/protein, Fig. S9c). For ESM2 and ProStT5, embedding generation times are lower: 12.59 ms and 76.55 ms, respectively. On the same mouse dataset executed using GPUs, FANTASIA v2 took a total of 7 h 38 min (6 h 10 min for the lookup and transfer step) while FANTASIA v1 took 1h 4 min 38 s. Memory usage also varies. FANTASIA v1’s RAM consumption scales linearly with the number of sequences (y = 4.4 + 0.0021x GB, x being the number of sequences in the proteome, Fig. S9d), while v2 maintains a stable memory footprint, benefiting from a more structured storage approach.

**Figure S9:**
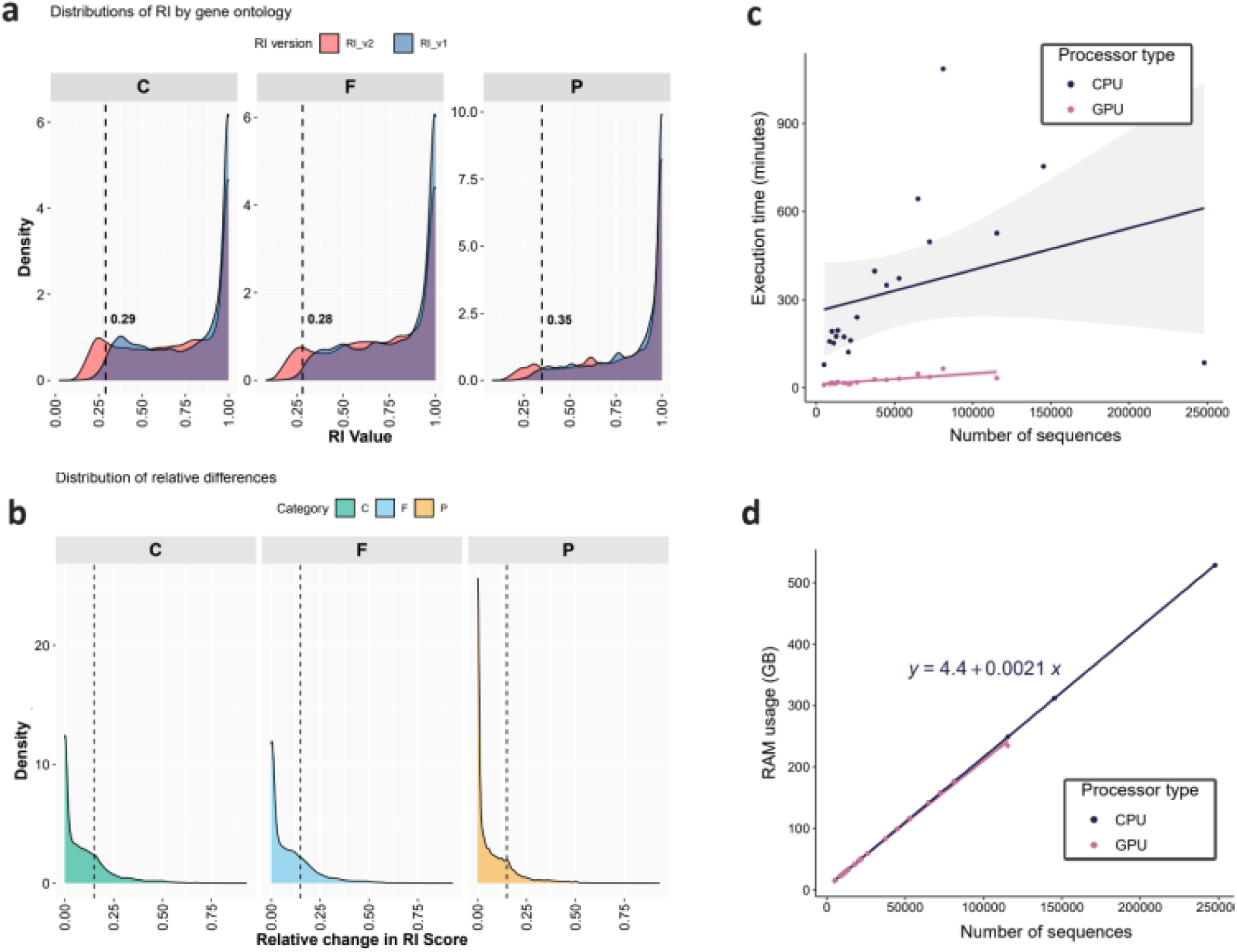
FANTASIA v1 vs v2. **a,** Distribution of RI-scaled Euclidean distances per ontology. The small discrepancies between both versions concentrate below the thresholds estimated in the GOPredSim publication^13^ (dashed vertical lines). **b,** Distribution of relative differences between both versions on 45,812 mouse proteins. The left of the dashed line represents the % of annotations having a difference of <15% (0.15 dashed line in the graph). These values are 85% for P, and around 75% for C and F. **c,** Relationship between the size of the proteome (number of sequences) and the execution time using CPUs or GPUs in FANTASIA v1. **d,** Relationship between the size of the proteome (number of sequences) and the memory employed when using CPUs or GPUs in FANTASIA v1.

## References

1. Ashburner, M. et al. Gene ontology: tool for the unification of biology. The Gene Ontology Consortium. Nat. Genet. 25, 25–29 (2000).

2. Cantalapiedra, C. P., Hernández-Plaza, A., Letunic, I., Bork, P. & Huerta-Cepas, J. eggNOG-mapper v2: Functional Annotation, Orthology Assignments, and Domain Prediction at the Metagenomic Scale. Mol. Biol. Evol. 38, 5825–5829 (2021).

3. Thomas, P. D. et al. PANTHER: Making genome-scale phylogenetics accessible to all. Protein Sci. 31, 8–22 (2022).

4. Barrios-Núñez, I., et al. Decoding functional proteome information in model organisms using protein language models. NAR Genom Bioinform 6, lqae078 (2024).

5. Hashimoto, T. et al. Extremotolerant tardigrade genome and improved radiotolerance of human cultured cells by tardigrade-unique protein. Nat. Commun. 7, 12808 (2016).

6. Kenny, N. J. et al. Tracing animal genomic evolution with the chromosomal-level assembly of the freshwater sponge Ephydatia muelleri. Nat. Commun. 11, 3676 (2020).

7. Lewin, H. A. et al. The Earth BioGenome Project 2020: Starting the clock. Proc. Natl. Acad. Sci. U. S. A. 119, (2022).

8. Mazzoni, C. J., Ciofi, C. & Waterhouse, R. M. Biodiversity: an atlas of European reference genomes. Nature 619, 252 (2023).

9. Mc Cartney, A. M., et al. The European Reference Genome Atlas: piloting a decentralised approach to equitable biodiversity genomics. bioRxiv 2023.09.25.559365 (2023) doi:10.1101/2023.09.25.559365.

10. Zhou, N. et al. The CAFA challenge reports improved protein function prediction and new functional annotations for hundreds of genes through experimental screens. Genome Biol. 20, 244 (2019).

11. Kulmanov, M. & Hoehndorf, R. DeepGOPlus: improved protein function prediction from sequence. Bioinformatics 36, 422–429 (2020).

12. Elnaggar, A. et al. ProtTrans: Toward Understanding the Language of Life Through Self-Supervised Learning. IEEE Trans. Pattern Anal. Mach. Intell. 44, 7112–7127 (2022).

13. Littmann, M., Heinzinger, M., Dallago, C., Olenyi, T. & Rost, B. Embeddings from deep learning transfer GO annotations beyond homology. Sci. Rep. 11, 1160 (2021).

14. Lin, Z. et al. Evolutionary-scale prediction of atomic-level protein structure with a language model. Science 379, 1123–1130 (2023).

15. Alexa, A. & Rahnenfuhrer, J. topGO. (Bioconductor, 2017). doi:10.18129/B9.BIOC.TOPGO.

16. Li, W. & Godzik, A. Cd-hit: a fast program for clustering and comparing large sets of protein or nucleotide sequences. Bioinformatics 22, 1658–1659 (2006).

17. Fu, L., Niu, B., Zhu, Z., Wu, S. & Li, W. CD-HIT: accelerated for clustering the next-generation sequencing data. Bioinformatics 28, 3150–3152 (2012).

18. Heinzinger, M., et al. Bilingual language model for protein sequence and structure. NAR Genomics and Bioinformatics 6, lqae150 (2024).

19. Pérez Canales, F. M. Fantasia Embeddings GOA2024. Zenodo 10.5281/zenodo.14864851 (2026).

20. PostgreSQL Global Development Group. pgvector: Open-source extension for vector similarity search in postgresql. pgvector https://github.com/pgvector/pgvector (2023).

21. Martínez-Redondo, G. I. et al. MATEdb2, a collection of high-quality metazoan proteomes across the animal tree of life to speed up phylogenomic studies. Genome Biol. Evol. 16, (2024).

22. Louie, B., Bergen, S., Higdon, R. & Kolker, E. Quantifying protein function specificity in the gene ontology. Stand. Genomic Sci. 2, 238–244 (2010).

23. Wang, J. Z., Du, Z., Payattakool, R., Yu, P. S. & Chen, C.-F. A new method to measure the semantic similarity of GO terms. Bioinformatics 23, 1274–1281 (2007).

24. Yoon, S., Baik, B., Park, T. & Nam, D. Powerful p-value combination methods to detect incomplete association. Sci. Rep. 11, 6980 (2021).

25. Sharma, N., Naorem, L. D., Jain, S. & Raghava, G. P. S. ToxinPred2: an improved method for predicting toxicity of proteins. Brief. Bioinform. 23, (2022).

26. Mirdita, M. et al. ColabFold: making protein folding accessible to all. Nat. Methods 19, 679–682 (2022).

27. Gu, Z. & Hübschmann, D. simplifyEnrichment: A Bioconductor Package for Clustering and Visualizing Functional Enrichment Results. Genomics Proteomics Bioinformatics 21, 190–202 (2023).

28. Klopfenstein, D. V. et al. GOATOOLS: A Python library for Gene Ontology analyses. Sci. Rep. 8, 10872 (2018).

29. Tomczak, A. et al. Interpretation of biological experiments changes with evolution of the Gene Ontology and its annotations. Sci. Rep. 8, 5115 (2018).

30. Martínez-Redondo, G. I. Data (part 1) from Illuminating the functional landscape of the dark proteome across the Animal Tree of Life through natural language processing models. Zenodo 10.5281/zenodo.10714961 (2024).

31. Martínez-Redondo, G. I. Data (part 2) from Illuminating the functional landscape of the dark proteome across the Animal Tree of Life through natural language processing models. Zenodo 10.5281/zenodo.10717484 (2024).

32. Martínez-Redondo, G. I. Data (part 3) from Illuminating the functional landscape of the dark proteome across the Animal Tree of Life through natural language processing models. Zenodo 10.5281/zenodo.10717774 (2024).

33. Martínez-Redondo, G. I. Data (part 4) from Illuminating the functional landscape of the dark proteome across the Animal Tree of Life through natural language processing models. Zenodo 10.5281/zenodo.10717783 (2024).

34. Martínez-Redondo, G. I. Data (part 5) from Illuminating the functional landscape of the dark proteome across the Animal Tree of Life through natural language processing models. Zenodo 10.5281/zenodo.10717885 (2024).

35. Martínez-Redondo, G. I. Data (part 6) from Illuminating the functional landscape of the dark proteome across the Animal Tree of Life through natural language processing models. Zenodo 10.5281/zenodo.10717910 (2024).

36. Thomas, P. D. The Gene Ontology and the Meaning of Biological Function. Methods Mol. Biol. 1446, 15–24 (2017).

37. Kober, K. M. & Pogson, G. H. Genome-wide signals of positive selection in strongylocentrotid sea urchins. BMC Genomics 18, 555 (2017).

38. McTaggart, S. J., Obbard, D. J., Conlon, C. & Little, T. J. Immune genes undergo more adaptive evolution than non-immune system genes in Daphnia pulex. BMC Evol. Biol. 12, 63 (2012).

39. Shultz, A. J. & Sackton, T. B. Immune genes are hotspots of shared positive selection across birds and mammals. Elife 8, (2019).

40. van Thiel, J. et al. Convergent evolution of toxin resistance in animals. Biol. Rev. Camb. Philos. Soc. 97, 1823–1843 (2022).

41. Kawabata, S. & Iwanaga, S. Role of lectins in the innate immunity of horseshoe crab. Dev. Comp. Immunol. 23, 391–400 (1999).

42. Carlson, D. E. & Hedin, M. Comparative transcriptomics of Entelegyne spiders (Araneae, Entelegynae), with emphasis on molecular evolution of orphan genes. PLoS One 12, e0174102 (2017).

43. Arzu Öztürk-Çolak, Steven J Marygold, Giulia Antonazzo, Helen Attrill, Damien Goutte-Gattat, Victoria K Jenkins, Beverley B Matthews, Gillian Millburn, Gilberto dos Santos, Christopher J Tabone, FlyBase Consortium. FlyBase: updates to the Drosophila genes and genomes database. Genetics 227, iyad211 (2024).

44. Kristensen, R. M. & Funch, P. Micrognathozoa: a new class with complicated jaws like those of Rotifera and Gnathostomulida. J. Morphol. 246, 1–49 (2000).

45. Li, Y., Shen, X.-X., Evans, B., Dunn, C. W. & Rokas, A. Rooting the Animal Tree of Life. Mol. Biol. Evol. 38, 4322–4333 (2021).

46. Burkhardt, P. et al. Syncytial nerve net in a ctenophore adds insights on the evolution of nervous systems. Science 380, 293–297 (2023).

47. Dallago, C., et al. Learned Embeddings from Deep Learning to Visualize and Predict Protein Sets. Curr Protoc 1, e113 (2021).

